# Microglial SIRPα regulates the emergence of CD11c^+^ microglia and demyelination damage in white matter

**DOI:** 10.1101/443531

**Authors:** Miho Sato-Hashimoto, Tomomi Nozu, Riho Toriba, Ayano Horikoshi, Miho Akaike, Kyoko Kawamoto, Ayaka Hirose, Yuriko Hayashi, Hiromi Nagai, Wakana Shimizu, Ayaka Saiki, Tatsuya Ishikawa, Ruwaida Elhanbaly, Takenori Kotani, Yoji Murata, Yasuyuki Saito, Masae Naruse, Koji Shibasaki, Pre-Arne Oldenborg, Steffen Jung, Takashi Matozaki, Yugo Fukazawa, Hiroshi Ohnishi

**Affiliations:** Department of Laboratory Sciences, Gunma University Graduate School of Health Sciences, Gunma 371-8514, Japan; Division of Brain Structure and Function, Life Science Innovation Center, Research Center for Child Mental Development, Faculty of Medical Sciences, University of Fukui, Fukui 910-1193, Japan; Department of Anatomy, Histology and Embryology, Faculty of Veterinary Medicine, Assiut University, Assiut-71526, Egypt; Division of Molecular and Cellular Signaling, Department of Biochemistry and Molecular Biology, Kobe University Graduate School of Medicine, Kobe 650-0017, Japan; Department of Molecular and Cellular Neurobiology, Gunma University Graduate School of Medicine, Gunma 371-8511, Japan; Department of Integrative Medical Biology, Section for Histology and Cell Biology, Umeå University, SE-901 87 Umeå, Sweden; Department of Immunology, The Weizmann Institute of Science, Rehovot 76100, Israel

**Author notes:** Corresponding author (H. Ohnishi).

## Abstract

A characteristic subset of microglia expressing CD11c appears in response to brain damage. However, the functional role of CD11c^+^ microglia, as well as the mechanism of its induction, are poorly understood. Here we report that the genetic ablation of signal regulatory protein α (SIRPα), a membrane protein, induced CD11c^+^ microglia in the brain white matter. Mice lacking CD47, a physiological ligand of SIRPα, and microglia-specific SIRPα knockout mice exhibited the same phenotype, suggesting the interaction between microglial SIRPα and CD47 on neighbouring cells suppressed the emergence of CD11c^+^ microglia. A lack of SIRPα did not cause detectable damage in the white matter, but resulted in the increased expression of genes characteristic of the repair phase after demyelination. In addition, cuprizone-induced demyelination was alleviated by the microglia-specific ablation of SIRPα. Thus, microglial SIRPα suppresses the induction of CD11c^+^ microglia that have the potential to accelerate the repair of damaged white matter.

## Introduction

Microglia constantly survey the microenvironment of the brain. When microglia encounter tissue damage, they become activated, produce multiple humoral factors and show enhanced phagocytic activity; microglia thus play a central role in the removal and repair of damaged tissues (Kettenmann, Hanisch, Noda, & Verkhratsky, 2011). However, the excessive activation of microglia results in the progression of inflammation and tissue degeneration, which is harmful to the brain environment (Kettenmann et al., 2011). The activation of microglia has been reported in various neurodegenerative diseases and brain injuries, including Alzheimer’s disease (AD), amyotrophic lateral sclerosis (ALS), and demyelinating diseases (Chiu et al., 2013; Holtman et al., 2015; Kamphuis, Kooijman, Schetters, Orre, & Hol, 2016; Remington, Babcock, Zehntner, & Owens, 2007; Wang et al., 2015). In animal models of these disorders, a characteristic subset of microglia that express integrin αX or CD11c, a marker of murine peripheral dendritic cells and selected macrophages, appear in the affected brain regions. The expression of CD11c is a marker for “primed” microglia that are not fully activated but rather are in a pre-activation state (Holtman et al., 2015; Norden & Godbout, 2013). CD11c^+^ microglia were also reported during postnatal development and normal aging (Bulloch et al., 2008; Kaunzner et al., 2012). In both cases, CD11c^+^ microglia characteristically appeared in the brain white matter, where myelin construction during the late developmental stage (Baumann & Pham-Dinh, 2001) or accumulation of damaged myelin debris during aging were remarkable (Safaiyan et al., 2016). It was also shown that demyelination markedly induced CD11c^+^ microglia, even in the adult brain (Remington et al., 2007), which was suppressed in mutant mice lacking Trem2 or Cx3Cr1 (Lampron et al., 2015; Poliani et al., 2015), i.e. functional molecules that promote phagocytosis. Of note, the clearance of myelin debris was markedly impaired in these mutant mice (Cantoni et al., 2015; Lampron et al., 2015; Poliani et al., 2015). CD11c^+^ microglia also accumulate around amyloid plaques in an AD mouse model with gene expression patterns indicating an enhanced capacity for phagocytosis (Kamphuis et al., 2016; Wang et al., 2015). Phagocytic ability of CD11c^+^ microglia thus might contribute to the removal and repair of damaged tissues. However, the physiological roles of CD11c^+^ microglia, as well as the mechanisms controlling the induction of CD11c^+^ microglia, are not fully understood.

SIRPα (CD172a) is a membrane protein highly expressed in macrophages and dendritic cells. The extracellular region of SIRPα specifically interacts with another membrane protein CD47 to promote cell–cell contact signals (Matozaki, Murata, Okazawa, & Ohnishi, 2009). Interactions between SIRPα on phagocytic cells and CD47 on phagocytosed targets, such as erythrocytes, cancer cells, and apoptotic cells, act as a “don’t eat me” signal and thereby control erythrocyte homeostasis, elimination of cancer cells, and the formation of arteriosclerotic plaques (Chao et al., 2011; Ishikawa-Sekigami et al., 2006; Kojima et al., 2016; Willingham et al., 2012). In the brain, SIRPα and CD47 are abundantly expressed on neurons (Ohnishi et al., 2010), and SIRPα is also expressed on microglia (Gitik, Liraz-Zaltsman, Oldenborg, Reichert, & Rotshenker, 2011). It is assumed that SIRPα on microglia interacts with neuronal CD47 to suppress microglial activation (Saijo & Glass, 2011), because SIRPα contains an immunoreceptor tyrosine-based inhibitory motif (ITIM) in its cytoplasmic region (Barclay & Hatherley, 2008; Matozaki et al., 2009). However, direct *in vivo* evidence supporting this model is missing. Here, we analysed the role of SIRPα in the control of microglial activation using CD47-SIRPα signal-deficient mice and define a critical microglia control module. Specifically, our results establish that SIRPα on microglia controls cell activation through the interaction with its ligand CD47 and negatively regulates the induction of CD11c^+^ microglia in the white, but not grey matter, which have the potential to support the repair of damaged myelin.

## Results

### Emergence of CD11c^+^ microglia in the brain white matter of SIRPα-deficient mice

To examine the role of SIRPα in microglial activation, brains of SIRPα knockout (KO) mice were subjected to immunohistochemical analysis using antibodies specific to Iba1, a microglia marker, and to CD68, a marker for phagocytically active microglia (Figure 1A). In the brains of SIRPα KO mice, numbers of Iba1^+^ as well as Iba1^+^/CD68^+^ cells were significantly increased in the white matter, such as the fimbria, compared with wild-type (WT) control mice, suggesting the activation of microglia in these regions (Figures 1A and 1B). Activation of microglia in the white matter was similar to the phenotype reported in aged mice, in which numbers of CD11c^+^ microglia were reported to be increased (Kaunzner et al., 2012). Next we examined the effect of the genetic ablation of SIRPα on the expression of CD11c on microglia by using CD11c-EYFP transgenic (Tg) mice (Lindquist et al., 2004), in which the expression of CD11c is detected with a reporter gene. In SIRPα KO:CD11c-EYFP Tg mice, Iba1^+^ cells and EYFP^+^ cells were markedly increased in the white matter, including the corpus callosum, external capsule, fimbria, and internal capsule, compared with control SIRPα^+/+^:CD11c-EYFP Tg mice (Figure 1C). Most EYFP^+^ cells in the white matter of SIRPα KO mice were Iba1 positive, suggesting they were CD11c^+^ microglia (Figure 1D ***lower panels***). In contrast, the EYFP signal was barely detected in the grey matter of both genotypes (Figure 1D ***upper panels***). The white matter-specific emergence of Iba1^+^/CD11c^+^ cells was also confirmed by using a specific antibody to CD11c in the brain or spinal cord of SIRPα KO mice (Figure 2A and 2B).

**Figure 1.**
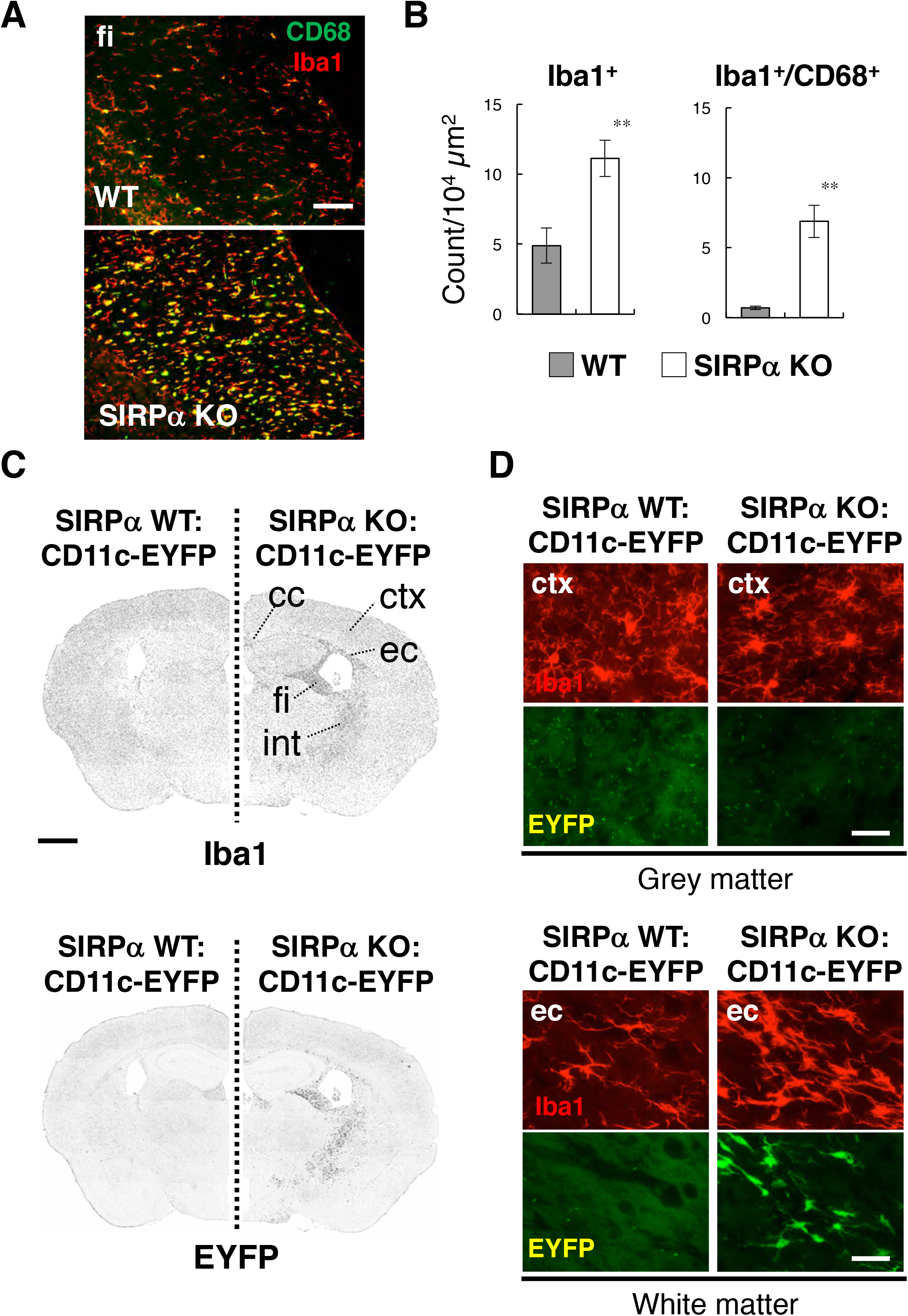
Activation of microglia in the brain white matter of SIRPα KO mice. (A) Immunofluorescence staining of coronal brain sections prepared from control (WT) or SIRPα null mutant mice (SIRPα KO) with antibodies to Iba1 (red) and CD68 (green). Merged images are shown. fi: fimbria. (B) Quantitative analysis of the number of Iba1^+^ (left panel) and Iba1^+^/CD68^+^ (right panel) microglia in the fimbria of WT (filled bars) and SIRPα KO mice (open bars) at 18–20 wks of age. Data are the means ± SEM (n = 6 images from 3 mice for each genotype). **P < 0.01 (Student’s t-test). (C, D) Immunofluorescence staining of coronal brain sections prepared from control CD11c-EYFP Tg mice carrying a homozygous SIRPα WT allele (SIRPα WT:CD11c-EYFP, left side) or SIRPα null mutation (SIRPα KO:CD11c-EYFP, right side) at 30–32 wks of age. Images in C are lower magnification images of the immunoreactivity of Iba1 (upper panels) and fluorescence of EYFP in the same sections (lower panels). Images were converted to grey scale and then inverted to clarify the fluorescence signal. Images in D are higher magnification images of the immunoreactivity of Iba1 (red) and fluorescence of EYFP (green) in grey (cerebrum cortex; ctx, upper panels) and white matter (external capsule; ec, lower panels). ctx: cerebrum cortex, cc: corpus callosum, ec: external capsule, fi: fimbria, int: internal capsule. Data are representative of at least 3 independent animals. Scale bars: 100 μm (A), 1 mm (C), and 50 μm (D).

**Figure 2.**
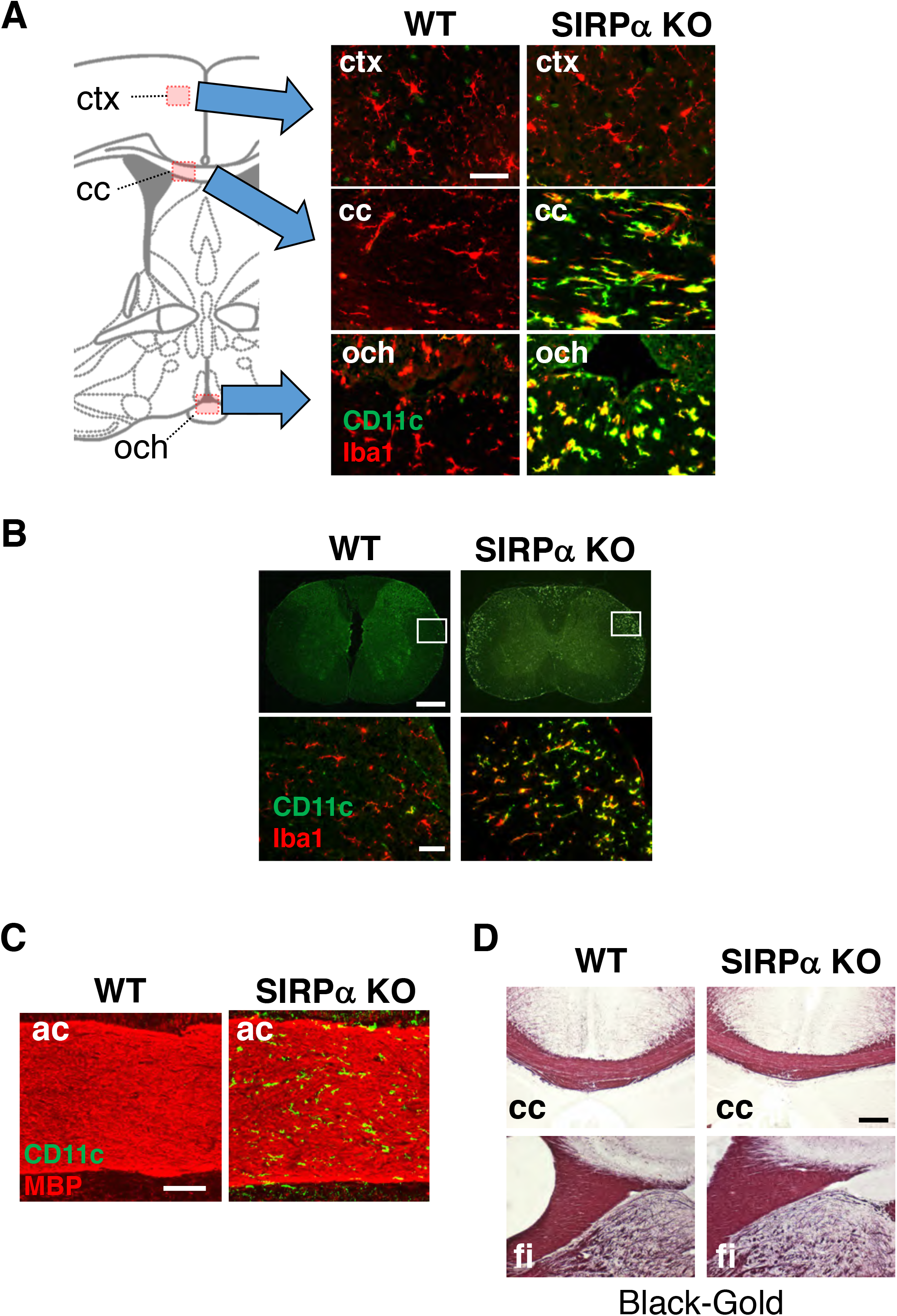
CD11c^+^ microglia in the white matter of SIRPα KO mice. (A, B) Coronal sections prepared from brain (A) and spinal cord (B) of control (WT) or SIRPα null mutant mice (SIRPα KO) at 18–20 wks of age with antibodies to Iba1 (red) and CD11c (green). In A, merged images are shown. Schematic diagrams of brain sections are shown on the left side. The boxed areas in B are shown at a higher magnification in the lower panels. ctx: cerebrum cortex, cc: corpus callosum, och: optic chiasma. (C, D) Coronal brain sections of control (WT) or SIRPα null mutant mice (SIRPα KO) at 29– 32 (C) or 10–11 (D) wks of age with antibodies to myelin basic protein (MBP) (red) and CD11c (green) (C), or Black-Gold (D). cc: corpus callosum, fi: fimbria, ac: anterior commissure. Data are representative of at least 3 independent animals. Scale bars, 50 μm (A, B lower panels), 1 mm (B upper panels), 100 μm (C), and 200 μm (D).

Demyelination damage causes microglia activation and an increase in the number of CD11c^+^ microglia in the white matter of the brain (Remington et al., 2007). Thus, we examined the structural integrity of myelin in SIRPα KO mice. Although a marked increase of CD11c^+^ cells was observed in the anterior commissure of SIRPα KO mice, no appreciable demyelination was evident by immunostaining for myelin basic protein (MBP) (Figure 2C). Myelination in the corpus callosum and fimbria were also normal in SIRPα KO mice when examined by myelin staining with a gold-phosphate complex (Black-Gold II) (Figure 2D).

### Increased expression of innate immune molecules in CD11c^+^ microglia in the brain of SIRPα-deficient mice and aged mice

We then examined the characteristics of the CD11c^+^ microglia. Microglia were isolated from the spinal cord of SIRPα KO and control WT mice, and the CD11b^+^/CD45^dim/lo^ fraction was analysed by flow cytometry (Figures 3A-3C). The yield of microglia from SIRPα KO mice was significantly higher than that from control mice [8.12 ± 0.47 and 4.57 ± 0.97 ×10^4^ cells/mouse, respectively; mean ± SEM (*n* = 5 animals), *P* = 0.01152 (Student’s *t*-test)], indicating an increased number of microglia in the spinal cord of SIRPα KO mice. As expected, expression of SIRPα was completely abolished in microglia prepared from SIRPα KO mice (Figure 3A). Consistent with the results from immunohistochemical analysis, CD11c^+^/CD11b^+^/CD45^dim/lo^ microglia were increased in SIRPα KO mice when compared with control WT mice (Figure 3B). Forward (FSC) and side (SSC) scatter distribution of CD11c^+^ microglia were similar to that of CD11c^—^ microglia in the SIRPα KO mice, suggesting cell size and granularity (complexity) were comparable between the two subsets (Figure 3—figure supplement 1). CD11c^+^ microglia expressed higher levels of innate immune molecules, including CD14, Dectin-1, and CD68 compared with CD11c^−^ microglia in SIRPα KO mice (Figure 3C). To compare CD11c^+^ microglia in SIRPα KO mice with those in aged WT mice (Bulloch et al., 2008; Kaunzner et al., 2012), microglia isolated from adult (16–18 weeks of age) and aged (69–105 weeks of age) WT mice were analysed in the same manner (Figure 3D-F). The expression levels of SIRPα in microglia were comparable between aged and adult mice (Figure 3D). Similar to SIRPα KO mice, CD11c^+^ microglia were increased in aged mice compared with adult mice (Figure 3E). In addition, CD11c^+^ microglia in the aged mice expressed higher levels of CD14, Dectin-1, and CD68 compared with CD11c^−^ microglia as in SIRPα KO mice (Figure 3F).

**Figure 3.**
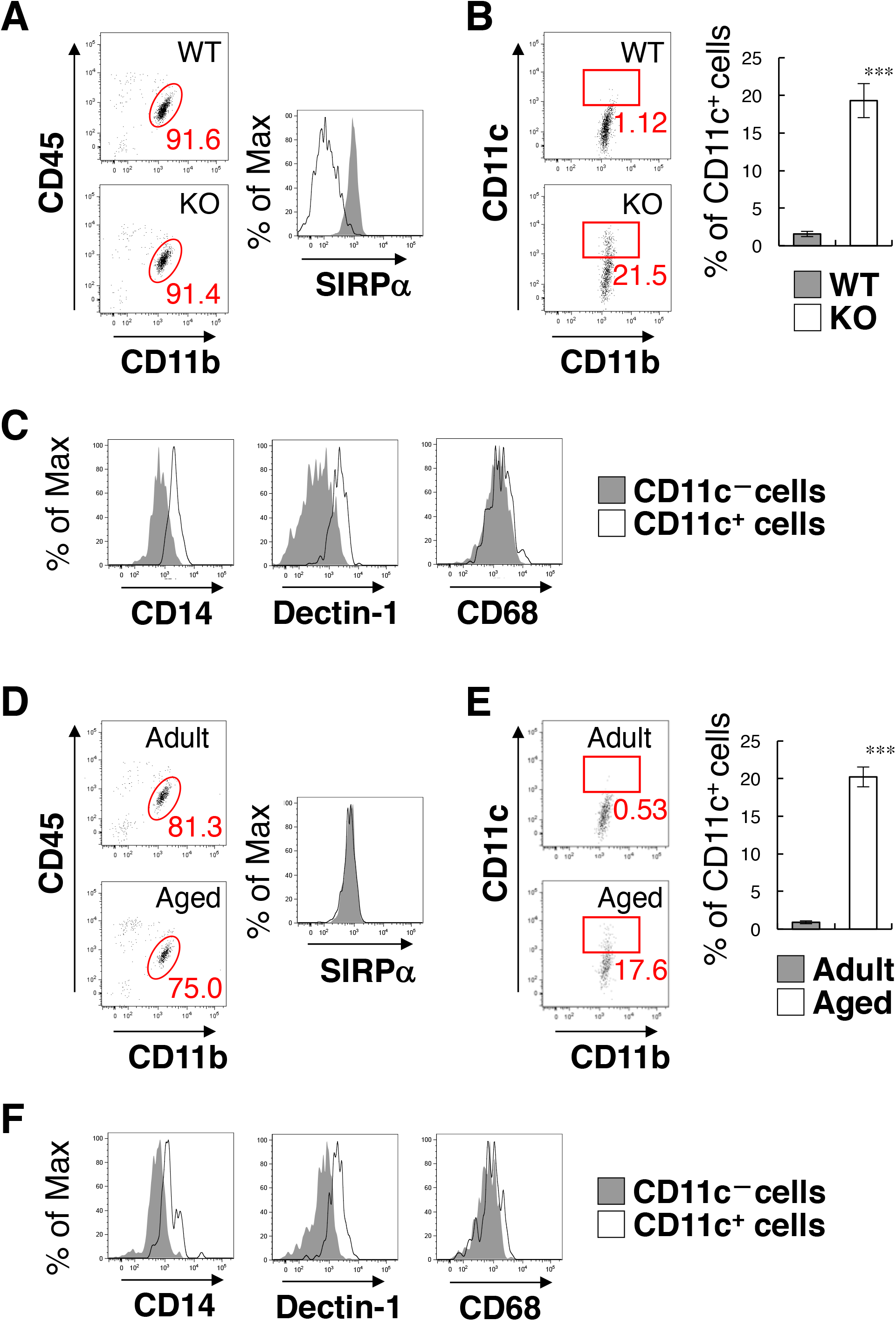
Flow cytometry analysis of microglia in SIRPα KO mice and WT aged mice. (A) Cells isolated from the spinal cord of control WT or SIRPα KO (KO) mice at 14 wks of age were incubated with a PE-conjugated mAb to SIRPα, a PerCP-Cy5.5– conjugated mAb to CD45, and an FITC-conjugated mAb to CD11b. The expressions of CD11b and CD45 on monocyte cells or of SIRPα on CD11b+/CD45^dim/lo^ microglia was analysed by flow cytometry. The percentage of CD11b+/CD45^dim/lo^ microglia among putative monocytes is indicated on each plot (left plots). Expression profiles for SIRPα on CD11b+/CD45^dim/lo^ microglia are shown in the right panel. (B) Cells prepared as in A from WT or SIRPα KO mice at 14–21 wks of age were stained with antibodies to CD45, CD11b, and CD11c, and analysed by flow cytometry. The percentage of CD11b+/CD45^dim/lo^/CD11c^+^ microglia among total CD11b+/CD45^dim/lo^ microglia is indicated on each plot. Quantitative data are shown in the right panel. (C) Cells prepared from SIRPα KO mice were stained for CD45, CD11b and CD11c, as well as CD14, Dectin-1, or CD68. Plots were gated on CD11b+/CD45^dim/lo^ cells and CD14, Dectin-1, or CD68 on CD11c-positive and –negative microglia were analysed. Expression profiles for each molecule in CD11b+/CD45^dim/lo^ microglia are shown. (D) Cells isolated from adult (18 wks of age) or aged (104 wks of age) mice were isolated and analysed as in A. (E) Cells prepared as in D from adult (16–18 wks of age) or aged (69–105 wks of age) mice were analysed as in B. Quantitative data are shown in the right panel. (F) Cells prepared from aged mice were analysed as in C. Data in B and E are the means ± SEM [n = 3 (B) and 5 (E) independent experiments]. ***P < 0.001 (Student’s t-test). Other data are representative of at least 3–5 independent experiments. Filled and open traces in (A), (C, F), and (D) indicate WT and SIRPα KO mice, CD11c— and CD11c^+^ cells, and adult and aged mice, respectively.

To address the effect of the increase of CD11c^+^ microglia on inflammatory responses, we examined the expression of pro- and anti-inflammatory cytokines in the brain and spinal cord by quantitative PCR analysis. Among the examined cytokines (TNF-α, IL-1β, IL-6, IL-10 and TGF-β), the expression of TNF-α was increased in SIRPα KO as compared to WT mice (Figure 3—figure supplement 2).

### Induction of CD11c^+^ microglia in the white matter of CD47-deficient mice

To address the mechanism involved in the regulation of microglia activation by SIRPα, we examined the effect of the genetic ablation of CD47, a membrane protein and SIRPα ligand (Matozaki et al., 2009). In the brain of CD47 KO mice, CD11c^+^ microglia were increased in the white matter as observed for the brains of SIRPα KO mice (Figure 4A). The number of Iba1^+^ microglia was increased more than 2.5-fold in the hippocampal fimbria, and about 70% of the Iba1^+^ microglia expressed CD11c (Figure 4B). Flow cytometric analysis showed that CD47 was expressed in WT microglia suggesting possible microglia-microglia communication through CD47-SIRPα interaction. Expression CD47 was completely absent in CD47 KO mice and the expression of SIRPα was markedly increased in CD47 KO microglia (Figure 4C). Similar to SIRPα KO mice, the yield of microglia from CD47 KO mice was increased compared with WT mice [7.65 ± 0.91 and 3.34 ± 0.84 ×10^4^ cells/head, respectively; mean ± SEM (*n* = 5 animals), *P* = 0.00866 (Student’s *t*-test)].

**Figure 4.**
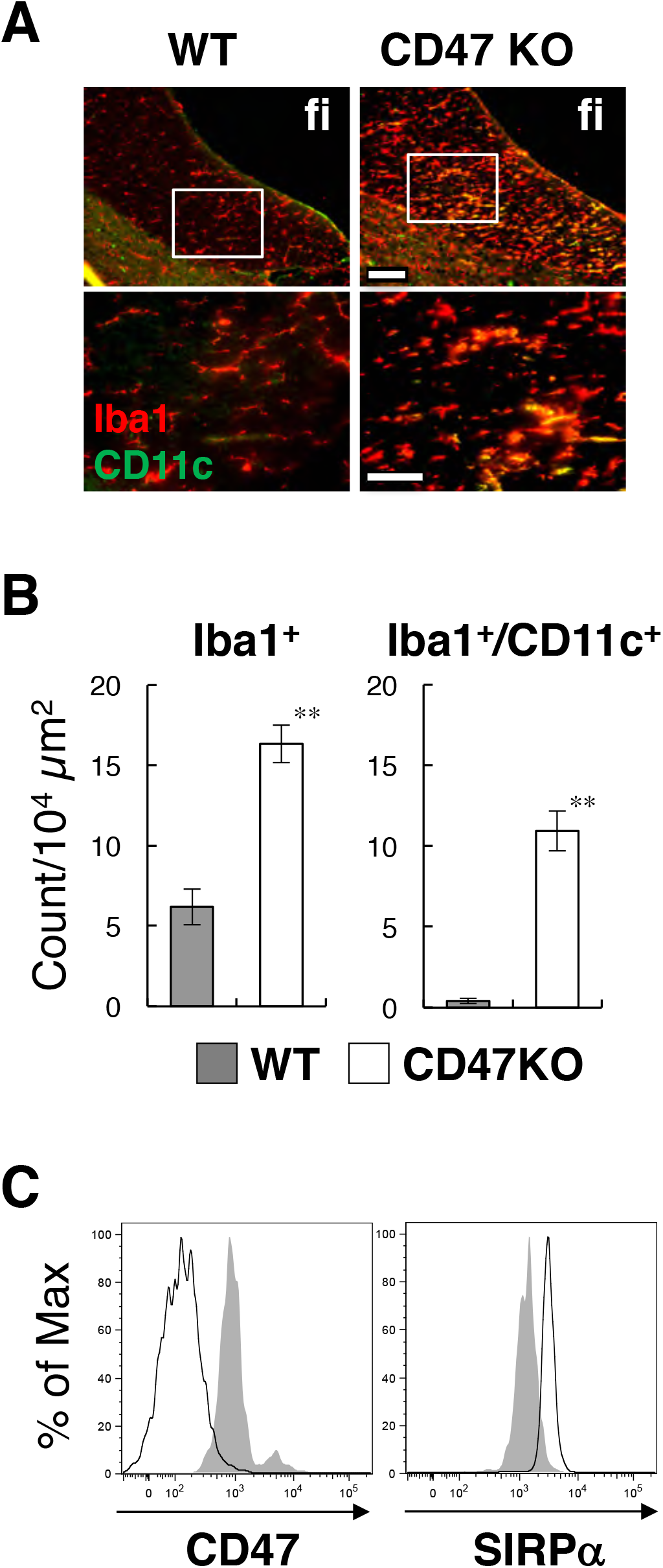
Activation of microglia in the brain white matter of CD47 KO mice. (A) Immunofluorescence staining of coronal brain sections prepared from control (WT) or CD47 KO mice at 19 wks of age with antibodies to Iba1 (red) and CD11c (green). Merged images are shown. The boxed areas in the upper panels are shown at higher magnification in the lower panels. fi: fimbria. Scale bars: 100 μm (upper panels), 50 μm (lower panels). (B) Quantitative analysis of the number of Iba1^+^ (left panel) and Iba1^+^/CD11c^+^ (right panel) microglia in the fimbria of WT (filled bars) and CD47 KO mice (open bars) at 13–27 wks of age. Data are the means ± SEM (n = 3 images from 3 mice for each genotype). **P < 0.01 (Student’s t-test). (C) Cells were isolated from the spinal cord of WT or CD47 KO mice at 14–16 wks of age. Expressions of CD47 or SIRPα on CD11b+/CD45^dim/lo^ microglia were analysed by flow cytometry. Data are representative of at least 3 independent experiments.

### Increased expression of genes for the repair of damaged myelin in CD47-deficient mice

To address the impact of the emergence of activated CD11c^+^ microglia on the brain environment, we compared the gene expression in the white matter (optic nerve and optic tract) between CD47 KO mice and WT control mice by microarray transcriptome analysis. Expression of total 14,875 genes was detected in both or either of the genotypes. Among them, 594 and 548 genes were markedly (> 2-fold) increased and decreased, respectively, in CD47 KO mice when compared with WT mice (**Supplementary file 1**). Pathway analysis with Database for Annotation, Visualization and Integrated Discovery (DAVID) ver. 6.8 (https://david.ncifcrf.gov/) (Huang, Sherman, & Lempicki, 2009a, 2009b) suggested that genes increased in the white matter of CD47 KO mice were significantly enriched in pathways such as infectious diseases, phagocytosis, and immune responses, likely reflecting the activation of microglia (Figure 5A). In contrast, genes decreased in CD47 KO mice were mostly enriched in pathways for neuronal ligand-receptor interaction, and were also enriched in pathways such as calcium signaling pathway and neuronal synapses (Figure 5B).

**Figure 5.**
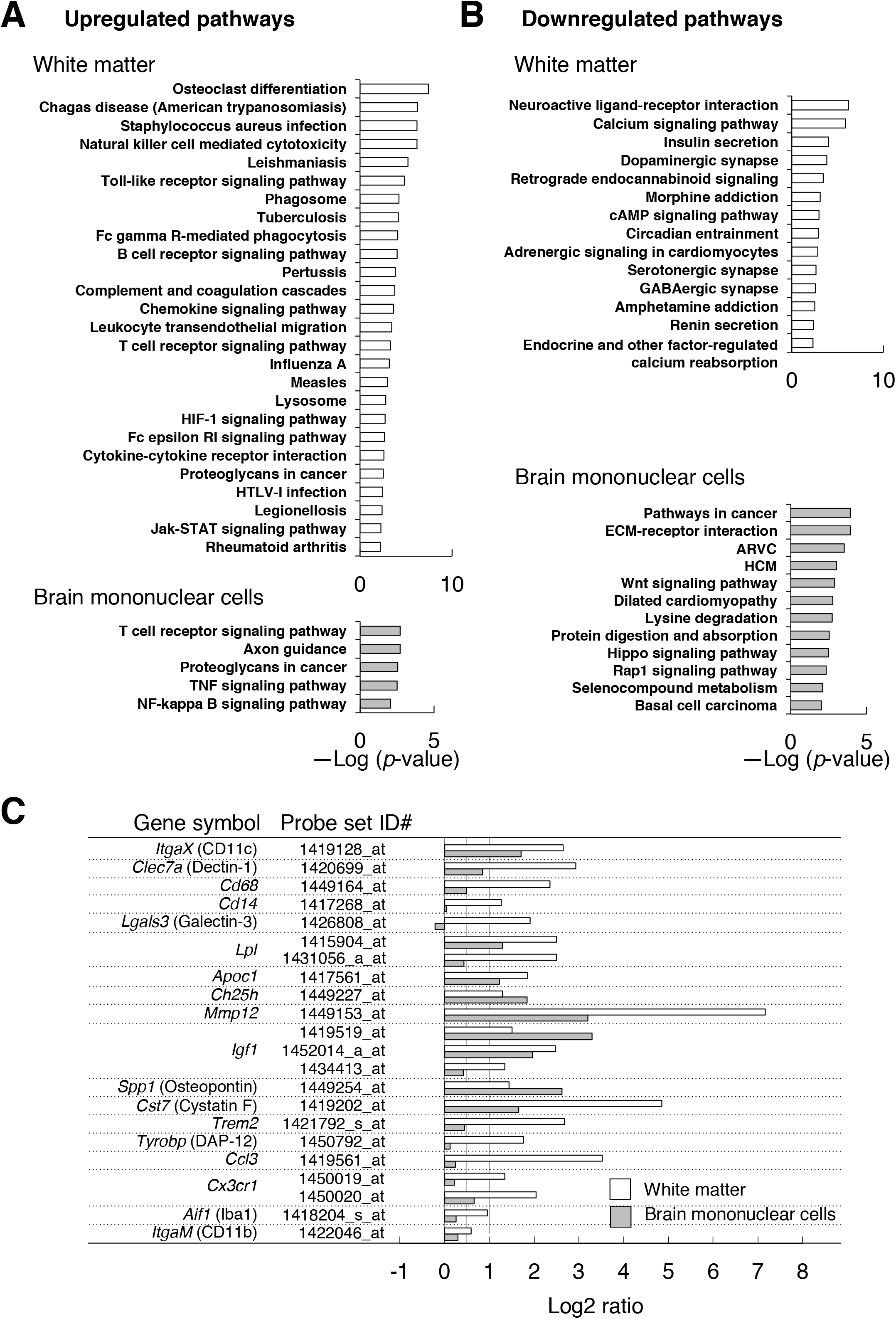
Microarray transcriptome analyses of the white matter and the brain mononuclear cells of CD47 KO mice. (A, B) The results of Kyoto Encyclopedia of Genes and Genomes (KEGG) pathway analysis with DAVID. Statistically significant (P-value < 0.01) KEGG enrichment pathways of up- (A) or down- (B) regulated genes in the white matter (optic nerve and optic tract) (upper panels) or in the brain mononuclear cells (lower panels) of CD47 KO mice were shown. Enrichment score is expressed as -Log (p-value). ARVC: Arrhythmogenic right ventricular cardiomyopathy, HCM: Hypertrophic cardiomyopathy. (C) The expression changes of the selected genes characteristics of microglia or of de- and re-myelination processes. For each probe set on the microarray, the fold-change of gene expression in the CD47 KO mice compared with WT mice was expressed as a log2 ratio. Open and grey bars show the comparison in the white matter and the brain mononuclear cells, respectively. Probe set ID#: Affymetrix probe set ID number.

Consistent with the results of immunohistochemical and flow cytometric analyses, the expression of *ItgaX* (CD11c), *Clec7a* (Dectin-1), *Cd68*, and *Cd14* were markedly increased (> 2-fold: Log2 ratio > 1) in the white matter of CD47 KO mice compared with WT mice (Figure 5C). In contrast, expression of the microglial markers *Aif1* (Iba1) and *ItgaM* (CD11b), was only moderately increased (< 2-fold). Thus, the marked increase in the expressions of CD11c and the other molecules was likely related to the characteristic change in the mutant microglia (or other cells) in the white matter, rather than to the increased number of microglia. Quantitative PCR analysis of selected genes (ItgaX, Igf1, Trem2, Ccl3), which were increased in the microarray analysis, also showed significant induction (Figure 5—figure supplement 1). We noted that several transcripts reported to be increased in microglia during the repair of damaged myelin (Holtman et al., 2015; Olah et al., 2012; Poliani et al., 2015; Selvaraju et al., 2004) were markedly increased (> 2-fold) in the white matter of CD47 KO mice (Figure 5C). These included *ItgaX, Lgals3*, and *Clec7a*, markers of primed microglia; *Lpl, Apoc1*, and *Ch25h*, molecules for lipid transport; *Mmp12*, a tissue remodelling factor; *Igf1*, and *Spp1*, trophic factors promoting oligodendrocyte differentiation; and *Cst7* (Cystatin F), a cysteine proteinase inhibitor. We also found that the positive regulators of microglia phagocytosis, *Trem2, DAP-12*, and *Cx3cr1* (Lampron et al., 2015; Poliani et al., 2015), were increased in the white matter of the mutant mice.

We next examined the gene expression in the brain mononuclear cells, in which microglia were enriched. Expression of total 16,544 genes was detected in both or either of CD47 KO and WT control cells. Among them, 1,323 and 2,286 genes were markedly (> 2-fold) increased and decreased, respectively, in the CD47 KO cells (**Supplementary file 2**). Genes increased in CD47 KO brain cells were significantly enriched in pathways for T cell receptor signal, axon guidance, proteoglycans in cancer, TNF signal, and NF-κ B signal (Figure 5A); genes decreased in CD47 KO brain cells were enriched in cancer associated pathways including Wnt, Hippo, and Rap1 signalling, as well as cardiomyopathy (Figure 5B).

Comparison of array data revealed 32 and 55 genes that were commonly increased and decreased (> 2-fold), respectively, in both the white matter and the brain mononuclear cells of CD47 KO mice (Figure 5—figure supplement 2). Shared induced genes included myelin-repair related genes, such as *ItgaX, Igf1, Lpl, Apoc1, Ch25h, Mmp12, Spp1*, and *Cst7* (Figure 5C). The expression of *Clec7a, CD68, Trem2*, and *Cx3cr1*, which were markedly increased in the white matter of CD47 KO mice, showed only moderate (< 2-fold) increase in the brain mononuclear cells (Figure 5C). Substantially higher expression of these genes in microglia might mask the increased expression of these genes in the limited CD11c^+^ subset that was only ~5 % of the total microglia when prepared from the whole brain of CD47 KO mice (data not shown).

### Induction of CD11c^+^ microglia in the brain white matter of microglia-specific SIRPα-deficient mice

To examine the cell type involved in the suppression of CD11c^+^ microglia by SIRPα, we analysed microglia-specific SIRPα conditional KO (cKO) mice that were generated by crossing SIRPα-flox mice (Washio et al., 2015) and Cx3cr1-CreER^T2^ mice to achieve microglia-specific gene targeting (Goldmann et al., 2013; Safaiyan et al., 2016; Wolf, Yona, Kim, & Jung, 2013). Flow cytometric analysis revealed that greater than 98% of microglia were SIRPα negative in the brain of tamoxifen-treated SIRPα^fl/fl^:Cx3cr1-CreER^T2^ mice (Figure 6A), and the numbers of Iba1^+^ and Iba1^+^/CD11c^+^ microglia were increased in the hippocampal fimbria of these cKO mice (Figures 6B and 6C). These data suggest that a lack of the interaction between SIRPα on microglia and CD47 on neighbouring cells is the primary cause for the induction of CD11c^+^ microglia in CD47-SIRPα signal-deficient mice. We also analyzed CD11c^+^ cell-specific SIRPα cKO mice (Washio et al., 2015). However, CD11c^+^ microglia were not increased in these mutant mice (Figure 6—figure supplement 1), suggesting that a lack of SIRPα in the resident CD11c^+^ microglia, a small subset of microglia in normal mouse brain (Bulloch et al., 2008), did not cause the expansion of these cells.

**Figure 6.**
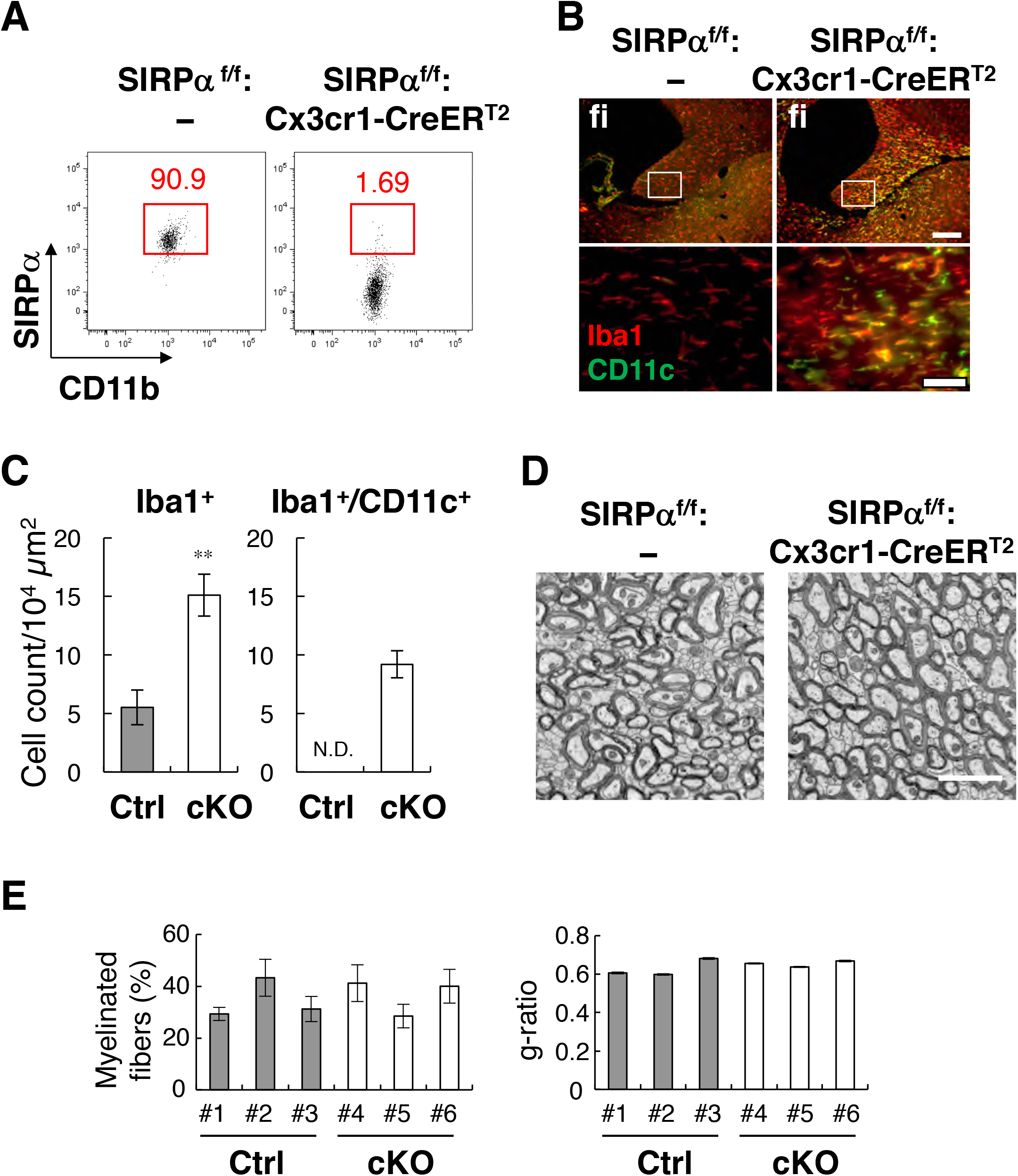
**Phenotypes of microglia-specific SIRPα conditional knockout mice**. (**A**) Cells isolated from the brains of control (**SIRPα^fl/fl^:—**) or microglia-specific SIRPα cKO (**SIRPα^fl/fl^:Cx3cr1-CreER^T2^**) mice at 11 wks of age were stained as in Figure 3A. The percentage of CD11b^+^/CD45^dim/lo^/SIRPα^+^ or /SIRPα^—^ microglia among total CD11b^+^/CD45^dim/lo^ microglia is indicated on each plot. Data are representative of at least 4 independent experiments. (**B**) Immunofluorescence staining of coronal brain sections prepared from control (SIRPα^fl/fl^) and SIRPα cKO mice at 22 wks of age with antibodies to Iba1 (***red***) and CD11c (***green***). Merged images are shown. The boxed areas in the upper panels are shown at higher magnification in the lower panels. **fi**: fimbria. Scale bars: 200 μm (***upper panels***), 50 μm (***lower panels***). (**C**) Quantitative analysis of the number of Iba1^+^ (***left panel***) and Iba1^+^/CD11c^+^ (***right panel***) microglia in the fimbria of control (SIRPα^fl/fl^) (**filled bars**) and SIRPα cKO (**cKO**) mice (**open bars**) at 22 wks of age. Data are the means ± SEM (*n* = 4 images from 2 mice for each genotype). \*\**P <* 0.01 (Student’s t-test). N.D.: not detected. (D) Electron microscopic analysis of axon fibres in the anterior commissure of control (SIRPα^fl/fl^:—) and microglia-specific SIRPα cKO mice (SIRPα^fl/fl^:Cx3cr1-CreER^T2^) at 26–27 wks of age. Scale bar: 2 μm. (E) Quantitative data of the myelination ratio (percentage of myelinated axons in total number of axons) and g-ratio (a ratio of the inner axonal diameter to the total outer diameter of myelinated axons) in the anterior commissure of three control (#1–#3, filled bars) and cKO mice (#4–#6, open bars). The myelination ratio and g-ratio were calculated from 9–10 images (109–465 axons/image) (n = 9–10) and 317–562 myelinated axons in 9–10 images (n = 317–562), respectively, for each mouse. Data are the means ± SEM.

### Alleviation of cuprizone-induced demyelination in the brain white matter of microglia-specific SIRPα-deficient mice

Although myelin damage induces CD11c^+^ microglia (Remington et al., 2007), demyelination was not observed in SIRPα total KO mice by light microscopy (Figures 2C and 2D). We examined the myelin structure again in microglia-specific cKO mice by electron microscopy (Figures 6D and 6E). In the cross section of the anterior commissure, the frequency of myelinated axons and the g-ratio (the ratio of the inner axonal diameter to the total outer diameter) were comparable between SIRPα cKO and control mice [frequency of myelinated axons: 34.3 ± 4.37% for control (n = 3), 36.3 ± 4.18% for cKO (n = 3), *P* = 0.757, Student’s t-test: g-ratio: 0.63 ± 0.025 for control (n = 3), 0.657 ± 0.009 for cKO (n = 3), *P* = 0.374, Student’s t-test], suggesting the myelin structure was normal in the mutant mice.

We further examined the progression of white matter damage in microglia-specific SIRPα cKO mice undergoing cuprizone (Cpz)-induced demyelination (Gudi, Gingele, Skripuletz, & Stangel, 2014; Norkute et al., 2009). Feeding with Cpz, a copper chelator, for 5 weeks (wks), induced demyelination accompanied by the robust accumulation of microglia (microgliosis) in white matter regions including the hippocampal alveus and corpus callosum in control mice as previously reported (Figure 7A) (Gudi et al., 2014; Norkute et al., 2009). At the site of microgliosis, CD11c^+^/Iba1^+^ microglia were markedly increased, even in control mice (Figure 7A). In SIRPα cKO mice, demyelination as well as microgliosis was significantly reduced after the same treatment (Figures 7A-7C). At an early stage of Cpz treatment (after 3 wks feeding with Cpz), demyelination was not observed in control or SIRPα cKO mice (Figure 7A), but abnormally strong immunoreactivity of MBP was observed (Figure 7A). This was probably related to myelin damage, because the epitope of the anti-MBP antibody we used (DENPVV) is similar to that of a myelin damage-detectable antibody (QDENPVV) (Matsuo et al., 1998). Consistently, significant microgliosis as well as an increase in CD11c^+^ cells were observed in both genotypes after 3 wks feeding with Cpz (Figures 7A and 7C). At this stage, the area of microgliosis was significantly larger in SIRPα cKO mice compared with control mice (Figures 7A and 7C). The repair of demyelination was observed after feeding with normal chow for 2 wks (Figure 7A). Microgliosis was still present in both genotypes at this time point but was significantly smaller in SIRPα cKO mice compared with control mice (Figures 7A and 7C). In the white matter, including the corpus callosum and alveus, the number of cells expressing Olig2+, an oligodendrocyte progenitor and lineage undergoing terminal differentiation into mature olidodendrocytes (Nishiyama, Komitova, Suzuki, & Zhu, 2009), was significantly decreased at 3 wks of Cpz feeding, suggesting demyelination was preceded by the loss of oligodendrocytes (Figures 7A and 7D). The number of Olig2^+^ cells was increased at 5 wks of Cpz treatment and recovered to normal levels after another 2 wks feeding with normal chow (Figures 7A and 7D). Throughout the de- and re-myelination process, no significant differences in the number of Olig2^+^ cells were noted between control and SIRPα cKO mice (Figures 7A and 7D).

**Figure 7.**
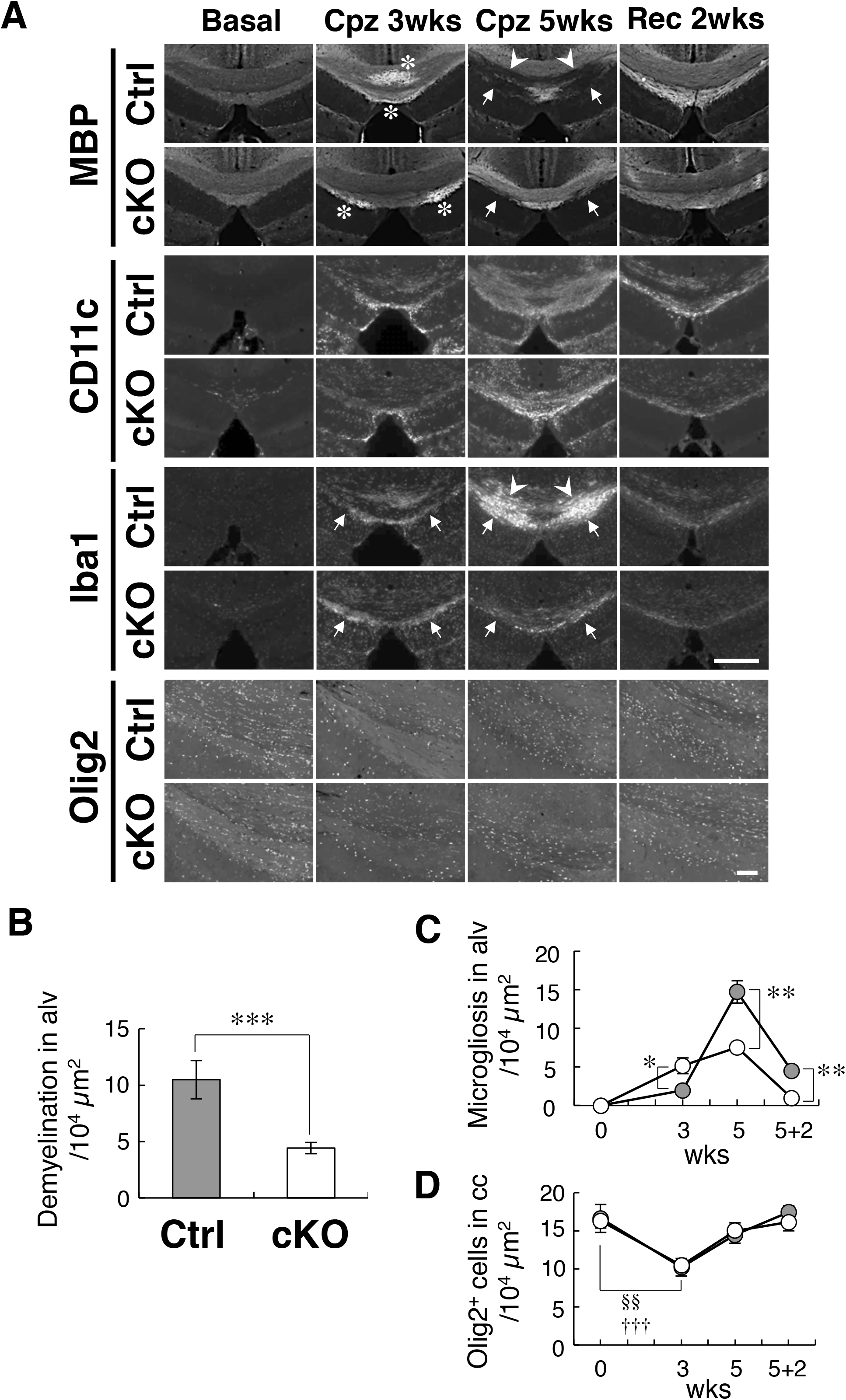
Alleviation of cuprizone-induced demyelination in SIRPα cKO mice. (A) Control SIRPα^fl/fl^ (Ctrl) or SIRPα cKO [SIRPα^fl/fl^:Cx3cr1-CreER^T2^ (cKO)] mice at 20– 25 wks of age were fed a 0.2% (w/w) cuprizone (Cpz) diet. After 3 or 5 wks of Cpz feeding (Cpz 3 wks, Cpz 5 wks), brain samples were prepared. Other groups of mice were returned to a normal diet after 5 wks of Cpz treatment and allowed to recover for 2 wks prior to the tissue analysis (Rec 2 wks). Mice fed a normal diet without Cpz for 7 wks were analysed as controls (Basal). Brain sections were subjected to immunofluorescence staining with specific antibodies for MBP, CD11c, Iba1, and Olig2. Arrowheads and arrows indicate typical demyelination (MBP) or microgliosis (Iba1) in the corpus callosum (arrowheads) and hippocampal alveus (arrows). Asterisks indicate abnormally strong immunoreactivity of MBP at Cpz 3 wks. Scale bars, 500 μm for MBP, CD11c, and Iba1, 100 μm for Olig2. (B, C) Areas with demyelination at Cpz 5 wks (B) and microgliosis at Basal, Cpz 3-5 wks, and Rec 2 wks (C) in the alveus (alv) were quantified by the immunoreactivity of MBP and Iba1, respectively. Data are the means ± SEM. A total of 8 areas in 4 images obtained from 4 mice (Cpz 5 wks), or a total of 4 areas in 2 images obtained from 2 mice (Basal, Cpz 3 wks, Rec 2 wks) were analysed for each genotype. *P < 0.05, **P < 0.01, ***P < 0.005 (Student‘s t-test). (D) The number of Olig2^+^ oligodendrocytes were quantified in the corpus callosum (cc). Data are the means ± SEM. A total of 10 images were obtained from 4 mice (Cpz 5 wks), or a total of 8 images were obtained from 2 mice (Basal, Cpz 3 wks, Rec 2 wks) and analysed for each genotype. §§P < 0.01 for Ctrl mice, †††P < 0.005 for cKO mice (Student’s t-test) versus the basal value of respective genotypes. No statistical significance was found between the two genotypes.

## Discussion

Cell–cell interactions between SIRPα on microglia and CD47 on neurons were proposed to suppress the activation of microglia (Ransohoff & Cardona, 2010). However, direct evidence for the function of this module in physiological context has not been provided. In this study, we demonstrated that SIRPα is a key molecule for the suppression of microglia activation *in vivo*. Molecules other than CD47, such as surfactant protein-A and -D (SP-A, -D), have also been reported as SIRPα ligands (Matozaki et al., 2009). However, we found that CD47 KO mice exhibited the same phenotype as SIRPα KO mice suggesting that the CD47-SIRPα interaction is indeed important for the suppression of microglia activation in the brain. In addition, microglia-specific SIRPα cKO mice also exhibited the same phenotype. Therefore, SIRPα expressed on microglia directly control microglia activation and the induction of CD11c^+^ microglia *in vivo*. The emergence of CD11c^+^ microglia in the brains of microglia-specific SIRPα cKO mice establishes that CD11c^+^ microglia are derived from resident microglia in the brain, and not recruited monocytes that are spared in the TAM-induced system (Goldmann et al., 2013; Safaiyan et al., 2016; Wolf et al., 2013). The characteristics of CD11c^+^ microglia in our mutant mice were similar to that of “primed microglia” that have been observed in aging and neurodegenerative diseases (Holtman et al., 2015); thus, SIRPα is a possible key component regulating microglia priming *in vivo*.

We found that the cell-surface expression of SIRPα was significantly increased in CD47 KO mice. It is likely that the binding of CD47 destabilises SIRPα molecules on microglia in the brain of WT mice, because our previous study suggested that the interaction between CD47 and SIRPα induced endocytosis of the CD47-SIRPα complex in CHO cells (Kusakari et al., 2008). It remains unclear which cell type expressed CD47 that contributed to the suppression of CD11c^+^ microglia. It was proposed that CD47 expressed on neurons suppresses the activation of microglia through interactions with SIRPα (Ransohoff & Cardona, 2010). An in vitro study suggested that CD47 on myelin sheets interacted with SIRPα and suppressed microglial phagocytosis (Gitik et al., 2011). Another study reported that direct cell–cell contact between astrocytes and microglia suppressed the expression of CD11c on microglia (Acevedo, Padala, Ni, & Jonakait, 2013). Thus, CD47 on oligodendrocytes or astrocytes may also participate in the suppression of microglia activation. Finally, we showed that microglia themselves express CD47 suggesting possible microglia-microglia interaction through CD47-SIRPα signal. Further research with cell type-specific CD47 conditional KO mice are required to clarify which cell type is involved in the CD47 mediated effects.

CD11c^+^ microglia are increased during neurodegenerative damage as well as in aging and postnatal development of normal brain (Bulloch et al., 2008; Kaunzner et al., 2012). We confirmed that CD11c^+^ microglia in SIRPα KO mice had elevated expressions of innate immune molecules including CD14, Dectin-1, and CD68, those are characteristic of CD11c^+^ microglia in WT aged mice. This suggested that the phenotype in the mutant mice was similar to normal biological responses in the aged brain. Although the maturation-dependent down regulation of CD47 was shown in hematopoietic stem cells (Jaiswal et al., 2009), the age-dependent reduction of CD47 has not been reported in the brain. In addition, our data suggested that the cell surface expression of SIRPα was not decreased in microglia from aged mice. Thus, the age-dependent dysfunction of CD47-SIRPα signals was unlikely. The increase of CD11c^+^ microglia in aged brains may be explained by the age-dependent accumulation of tissue damage that stimulates microglia activation beyond the suppressive ability of SIRPα.

We demonstrated that CD11c^+^ microglia were specifically increased in the white matter of mutant mice. Thus, the white matter may contain an endogenous factor that promotes the induction of CD11c^+^ microglia in mutant mice. One potential candidate is myelin. As reported (Poliani et al., 2015), CD11c^+^ microglia were increased in control SIRPα+/^+^ mice after Cpz treatment. Thus, myelin damage effectively induces CD11c^+^ microglia even in the control mice. This suggests that myelin degradation products during homeostatic turnover of the myelin structure might stimulate the induction of CD11c^+^ microglia in the CD47-SIRPα signal-deficient mice, while the effect of such homeostatic degradation of myelin may be suppressed by SIRPα in the normal mouse brain. Enhanced phagocytosis of myelin components by microglia may be involved in the induction of CD11c^+^ microglia in our mutant mice, because SIRPα negatively regulated phagocytosis in macrophages (Matozaki et al., 2009). CD11c^+^ microglia in SIRPα-deficient mice expressed CD68, a phagocytic marker, supporting the notion that phagocytic activity was increased in these cells. Several studies have suggested the importance of myelin phagocytosis by microglia for the induction of CD11c^+^ microglia during demyelination or aging. The myelin damage-dependent induction of CD11c^+^ microglia was markedly suppressed in mutant mice lacking Trem2, a phagocytic receptor (Poliani et al., 2015). The emergence of CD11c^+^ microglia during demyelination was also suppressed in Cx3cr1 KO mice, in which the phagocytosis of myelin debris by microglia was severely impeded (Lampron et al., 2015). In the brain of aged mice, the accumulation of myelin debris and phagocytosis of such components by microglia were increased in the white matter (Safaiyan et al., 2016), where CD11c^+^ microglia were characteristically observed in aged mice (Hart, Wyttenbach, Perry, & Teeling, 2012). Of note, the aging-induced expansion of microglia in the corpus callosum was suppressed in Trem2 KO mice (Poliani et al., 2015). Furthermore, an in vitro study suggested that SIRPα on microglia inhibited the phagocytosis of myelin membrane fractions through interactions with CD47 on myelin (Gitik et al., 2011). Thus, our present data support the model whereby the lack of CD47-SIRPα signal upregulates the phagocytosis of microglia, which become hypersensitive to the homeostatic destruction of myelin structures during the normal turnover process, and thereby triggers the induction of CD11c^+^ microglia without damage.

Transcriptome analyses showed that the expression of TNF-α, a proinflammatory cytokine, was increased 2-3-fold in the brain monocytes isolated from CD47 KO mice. In addition, pathway analysis showed other genes related to TNF and NF-κ B signalling pathway, including RelA and IKKB (*Ikbkb*), were specifically increased in the mutant brain cells. Thus, it is likely that proinflammatory TNF axis is strengthened in microglia by the lack of CD47-SIRPα signal. In contrast, Wnt signal pathway, which contributes to the proinflammatory transformation of microglia (Halleskog et al., 2011), was attenuated in the CD47 KO brain cells. The transcriptome analysis also showed that expression of TGF-β, an anti-inflammatory cytokine, and brain-derived neurotrophic factor (BDNF), a neuroprotective factor, were increased in the CD47 KO brain cells. Thus, it is not clear whether CD11c^+^ microglia in the mutant mice have pro or anti-inflammatory roles.

The expression of genes characteristic during the recovery phase from demyelination (Holtman et al., 2015; Olah et al., 2012; Poliani et al., 2015), such as *ItgaX, Igf1, Lpl, Apoc1, Ch25h, Mmp12, Spp1*, and *Cst7* were increased in both of the white matter and the brain cells of CD47 KO mice, suggesting their upregulation in the white matter CD11c^+^ microglia, and also suggesting the potential of the CD11c^+^ microglia to promote white matter tissue repair. Consistently, both demyelination and microgliosis after 5 wks of Cpz feeding were significantly reduced in SIRPα cKO mice. Although the direct involvement of CD11c^+^ microglia on the reduction of demyelination damage in the mutant mice is not yet clear, CD11c^+^ microglia are likely to have a protective effect against Cpz-induced demyelination. IGF-1 and Spp1 were increased in the white matter and brain mononuclear cells in CD47 KO mice. These factors derived from microglia and support the proliferation and differentiation of oligodendrocytes and myelination (Olah et al., 2012; Selvaraju et al., 2004; Zeger et al., 2007).

CD11c^+^ microglial in neonatal brains are reported as the major source of IGF-1 and to support primary myelination (Wlodarczyk et al., 2017). CD11c^+^ microglia in our mutant mice could facilitate white matter repair through the function of these factors. At early stages of Cpz treatment (3 wks feeding with Cpz), microgliosis was significantly more severe in SIRPα cKO mice compared with control mice, suggesting a faster response of SIRPα-deficient microglia to oligodendrocyte damage. Although Cpz feeding induced CD11c^+^ microglia in both control and SIRPα cKO mice, the preceding existence of CD11c^+^ microglia in SIRPα cKO mice may enable these cells to respond earlier to the damage and accelerate tissue repair by their protective effect.

The loss and recovery of Olig2^+^ cells were comparable between control and SIRPα cKO mice during de- and re-myelination. Cell death and restorative proliferation of the oligodendrocyte linage thus seems to occur in both groups to a similar extent. The alleviated demyelination in SIRPα cKO mice may be explained by the faster differentiation and/or maturation of oligodendrocytes compared with control mice, resulting in accelerated remyelination in the cKO mice. Another possibility is that both the cell death and restorative proliferation of oligodendrocytes were suppressed in SIRPα cKO mice, resulting in reduced damage compared with control mice. In this case, the total number of Olig2^+^ cells, the summation of surviving and proliferating Olig2^+^ cells, in SIRPα cKO would be comparable to control mice, in which both the loss and restoration of Olig2^+^ cells were much greater than in cKO mice, with greater demyelination damage.

Our data suggest a supportive role for CD11c^+^ microglia in tissue repair. Consistently, the suppression of the emergence of CD11c^+^ microglia by the genetic ablation of Trem2 was concurrent with exacerbated neuronal damage in demyelination model mice or neuronal death and amyloid deposition in AD model mice (Poliani et al., 2015; Wang et al., 2015). The emergence of CD11c^+^ microglia in aged brain might be an adaptive reaction to cope with tissue damage caused by aging. It is also notable that gene sets increased in the white matter of CD47 KO mice (i.e. ItgaX, lpl, Cst7, Clec7a, Trem2, and Tyrobp) were similar to those increased in DAM (disease-associated microglia), a potential protective microglia subtype associated with neurodegeneration such as AD (Keren-Shaul et al., 2017). CD47-SIRPα interactions may be a key mechanism for the induction of protective microglia such as DAM.

The reduction of demyelination in Cpz-treated SIRPα KO mice is in contrast to persistent demyelination after Cpz treatment in Trem2 KO mice (Poliani et al., 2015). In addition, the enhanced expression of *ItgaX, Igf1, Lpl, Ch25h*, and *Apoc1* in CD47 KO mice was markedly in contrast to that in Trem2 KO mice, in which the upregulation of these genes during demyelination in control mice were markedly suppressed (Poliani et al., 2015). The signalling mechanisms of SIRPα and Trem2 are contrasting. SIRPα activates tyrosine phosphatase Shp1/2, which binds to a phosphorylated ITIM in the cytoplasmic region of SIRPα, while Trem2 activates tyrosine kinase Syk, which binds to phosphorylated ITAM (immunoreceptor tyrosine-based activation motif) in the cytoplasmic part of DAP12, a membrane protein forming a receptor complex with Trem2 (Paradowska-Gorycka & Jurkowska, 2013). SIRPα and Trem2 may reciprocally control microglia phagocytosis through the function of tyrosine phosphatase and kinase, respectively. Further analysis of the interaction between CD47-SIRPα signalling and Trem2-DAP12 signalling may be important to understand microglial activation and phagocytosis and to identify new therapeutic strategies to promote tissue repair in the damaged brain.

### Experimental Procedures

#### Animals

SIRPα^—/—^ (SIRPα KO), CD47^—/—^ (CD47 KO), SIRPα^flox/flox^, and Cx3cr1-CreER^T2^ Tg mice were generated as described (Oldenborg et al., 2000; Washio et al., 2015; Yona et al., 2013). CD11c-Cre Tg mice [B6.Cg-Tg(Itgax-Cre)1-1Reiz/J, #008068] (Caton, Smith-Raska, & Reizis, 2007) and CD11c-EYFP transgenic mice [B6.Cg-Tg(Itgax-Venus)1Mnz/J, #008829] (Lindquist et al., 2004) were obtained from Jackson Laboratory (Bar Harbor, ME). SIRPα KO mice were crossed with CD11c-EYFP Tg mice to generate SIRPα^—/+^:CD11c-EYFP Tg mice. These mice were crossed with heterozygous SIRPα^—/+^ mice, and the resulting homozygous SIRPα WT and KO mice harbouring the CD11c-EYFP transgene were analysed. To obtain SIRPα cKO mice, homozygous SIRPα-floxed mice harbouring the CD11c-Cre or Cx3cr1-CreER^T2^ transgene were crossed with homozygous floxed mice, and the resulting SIRPα cKO (SIRPα^flox/flox^:Cx3cr1-CreER^T2^, SIRPα^flox/flox^:CD11c-Cre) mice were analysed. Littermates carrying homozygous SIRPα-flox alleles but lacking Cre recombinase were used as controls. To induce Cre-dependent recombination, Cx3cr1-CreER^T2^ mice were treated with tamoxifen (TAM) (Toronto Research Chemicals Inc., Ontario, Canada) at 8 wks of age. TAM was dissolved in ethanol and then diluted 7 times with corn oil (Wako, Osaka, Japan) to make a 10 mg/ml solution, and 200 μL (2 mg) of this solution was injected subcutaneously once every 48 h for 5 consecutive days (total of 3 times). TAM-treated mice were analysed more than 8 wks after the treatment. All mice were bred and maintained at the Bioresource Center of Gunma University Graduate School of Medicine under specific pathogen–free conditions. Mice were housed in an air-conditioned room with a 12-h-light, 12-h-dark cycle. All animal experiments were approved by the Animal Care and Experimentation Committee of Gunma University (approval no. 13-009).

#### Primary antibodies and reagents

Rabbit polyclonal antibodies (pAbs) to Iba1 were obtained from Wako (Osaka, Japan). Biotin-conjugated Armenian Hamster monoclonal antibody (mAb) to mouse CD11c (clone HL3), PerCP-Cy5.5 conjugated rat mAb to CD45 (30-F11), FITC-conjugated rat mAb to mouse CD11b (clone M1/70), rat mAb to mouse CD16/CD32 (clone 2.4G2), and streptavidin-conjugated allophycocyanin (APC) were from BD Pharmingen (San Diego, CA). PE conjugated rat mAb to mouse CD172a (SIRPα) (clone P84) was from eBioscience (San Diego, CA). PE conjugated and unconjugated rat mAbs to CD68 (clone FA-11) and PE conjugated rat mAb to mouse CD14 (clone Sa14-2) were from BioLegend (San Diego, CA). PE conjugated recombinant antibody (Ab) to Dectin-1 (REA154) and PE conjugated recombinant Ab to CD47 (REA170) were from Miltenyi Biotec (Bergisch Gladbach, Germany). 4′,6-Diamidino-2-phenylindole (DAPI) was obtained from Nacalai Tesque (Kyoto, Japan). Rat mAb to myelin basic protein (MBP) (clone 12) was from Merck Millipore (Billerica, MA). Rabbit pAb to Olig2 (18953) was from Immuno-Biological Laboratories (Minneapolis, MN).

#### Histological analysis

For immunohistochemistry, mice were anesthetised by the inhalation of isoflurane (Pfizer, New York, NY) and an intraperitoneal injection of pentobarbital (Nembutal 100 mg/kg; Dynabot, Tokyo, Japan), and then perfused transcardially with fixation buffer [4% paraformaldehyde in 0.1 M phosphate buffer (pH 7.4)]. Brain or spinal cord was dissected and fixed in the fixation buffer overnight at 4°C, then transferred to 30% sucrose in 0.1 M phosphate buffer (pH 7.4) for cryoprotection, embedded in OCT compound (Sakura Fine Technical, Tokyo, Japan), and rapidly frozen in liquid nitrogen. Frozen sections with a thickness of 10 or 20 μm were prepared with a cryostat, mounted on glass slides, air dried, and washed with phosphate buffered saline (PBS). Otherwise, free floating sections were washed with PBS. All sections were then incubated for 30 min at room temperature in Tyramide Signal Amplification (TSA) blocking solution (Perkin Elmer, Norwalk, CT) and stained overnight at 8°C with primary antibodies diluted in primary antibody dilution buffer (PBS supplemented with 2.5% BSA and 0.3% Triton X-100). Sections were then washed with PBS and exposed to the corresponding secondary antibodies conjugated with the fluorescent dyes Cy3 (Jackson Immuno Research Laboratories, West Grove, PA) or Alexa Fluor 488 (Invitrogen, Carlsbad, CA) in secondary antibody dilution buffer (PBS supplemented with 1% BSA and 0.3% Triton X-100). Nuclei were also stained with DAPI. The detection of CD11c with biotin conjugated primary antibody (HL3) was achieved by using a TSA Biotin System kit (Perkin Elmer) and streptavidin-conjugated Alexa Fluor 488 (Invitrogen) according to the manufacturer’s protocol. Free floating sections were mounted on glass slides after staining. All sections were mounted with mounting media [0.1 M Tris-HCl buffer (pH9) containing 5% 1,4-diazabicylo [2.2.2]-octane (DABCO), 20% polyvinyl alcohol (PVA), 10% glycerol] and covered with a cover glass. Fluorescence images were acquired with a fluorescence microscope BZ-X710 (Keyence, Osaka, Japan) or DM RXA (Leica, Wetzlar, Germany) equipped with a cooled CCD camera (Cool SNAP HQ; Roper Scientific, Trenton, NJ). Acquired digital images were analysed by ImageJ software (Schneider, Rasband, & Eliceiri, 2012).

For myelin staining, frozen brain sections were mounted on a glass slide first, and then stained with a Black-Gold II myelin staining kit (Merck Millipore) according to the manufacturer’s protocol. Images were acquired with a light microscope DM IRBE (Leica) equipped with a cooled CCD camera (Penguin 600CL; Pixera Corp., Santa Clara, CA).

#### Preparation of microglia and flow cytometry

Mononuclear cells including microglia were isolated by the method described by Sierra et al. (Sierra, Gottfried-Blackmore, McEwen, & Bulloch, 2007) with a minor modification. Briefly, brain or spinal cord was dissected from genotype- and age-matched male mice euthanised by cervical dislocation after deep anaesthesia. Spinal cords isolated from 1–3 mice were mixed and treated together. Tissues were rinsed in ice-cold Hank’s Balanced Salt Solution (HBSS; Gibco, Grand Island, NY) and then homogenised in HBSS containing collagenase type 2 (37.5 U/ml; Worthington) and DNase I (45 U/ml; Sigma). Homogenates were then incubated at 37°C for 30 min, gently triturated with a Pasteur pipette, and incubated at 37°C for 10 min. Undigested material was removed by filtration through a 70-μm cell strainer (BD Biosciences), and the remaining cells were centrifuged at 1,100 ×g for 5 min at 18°C. Collected cells were then suspended in 8 ml of 1×HBSS containing 70% Percoll (GE Healthcare Life Science, Uppsala, Sweden), and then 4–5 ml of each was placed into a 15 ml tube. Cell suspensions were then overlaid consecutively with 4 ml HBSS containing 37% Percoll and 1 ml PBS. The resulting gradient was centrifuged at 200 ×g for 40 min at 18°C, after which cells at the interface of the bottom 2 layers (70%/37%) were collected, washed twice with HBSS, and subjected to flow cytometric analysis.

For flow cytometry analysis, cells were first incubated with a mAb specific for mouse CD16/CD32 to prevent nonspecific binding of labelled mAbs against FcγR and were then labelled with specific mAbs conjugated with PE, FITC, PerCP-CyTM5.5, or biotin. For the staining of CD11c, a biotin-conjugated mAb against mouse CD11c was detected with streptavidin-conjugated APC. Labelled cells were analysed by flow cytometry using a BD FACS Canto II flow cytometer (BD Biosciences, San Jose, CA). Cells were first gated on their forward (FSC) and side (SSC) scatter properties to discriminate putative monocytes from other events, and then were gated on CD45 and CD11b. The CD11b+/CD45^dim/lo^ fraction obtained was analysed as microglia. All data were analysed with FlowJo 8.8.4 software (Tree Star Inc., Ashland, OR).

#### Quantitative PCR analysis

Total RNA was extracted from the whole brain, spinal cord, or dissected optic nerve and optic tract with the use of Sepasol RNA I (Nacalai Tesque) and an RNeasy Mini kit (Qiagen, Hilden, Germany). First-strand cDNA was synthesised from total RNA with the use of a QuantiTect Reverse Transcription kit (Qiagen), and cDNA fragments of interest were amplified by real-time PCR in 96-well plates (Roche Diagnostics, Mannheim, Germany) with the use of a QuantiTect SYBR Green PCR kit (Qiagen) or FastStart SYBR Green Master (Roche Diagnostics, Mannheim, Germany) and LightCycler 480 or 96 Real-Time PCR System (Roche Diagnostics). The amplification results were analysed with the use of LightCycler software and were then normalised on the basis of the *Gapdh* mRNA level in each sample. Primer sequences (forward and reverse, respectively) were as follows: *Tnfa*, 5‘-CCCTCACACACTCAGATCATCTTCT-3’ and 5’-GCTACGACGTGGGCTACAG-3’; *Il1b*, 5’-CAACCAACAAGTGATATTCTCCATG-3’ and 5’-GATCCACACTCTCCAGCTGCA-3’; *Il6*, 5’-TAGTCCTTCCTACCCCAATTTCC-3’ and 5‘-TTGGTCCTTAGCCACTCCTTC-3’; *Il10*, 5’-AGGCGCTGTCATCGATTTCT-3’ and 5’-ATGGCCTTGTAGACACCTTGG-3’; *Tgfb*, 5‘- ACCATGCCAACTTCTGTCTG-3’ and 5’-CGGGTTGTGTTGGTTGTAGA-3’; *ItgaX*, 5’-AGCTGTGTGGACAGTGATGG-3’ and 5‘-TGCATGTGAGTCAGGAGGTC-3’; *Igf1*, 5’-TACTTCAACAAGCCCACAGGC-3’ and 5’-ATAGAGCGGGCTGCTTTTGT-3’; *Trem2*, 5‘-CTTCCTGAAGAAGCGGAATG -3’ and 5’-AGAGTGATGGTGACGGTTCC-3’; *Ccl3*, 5’-CAGCCAGGTGTCATTTTCCT-3’ and 5’-CTGCCTCCAAGACTCTCAGG -3’; and *Gapdh*, 5’-TCCCACTCTTCCACCTTCGA-3’ and 5’-GTCCACCACCCTGTTGCTGTA-3’.

#### Microarray analysis

Total RNAs were prepared from the white matter (optic nerve and optic tract) or brain mononuclear cells of WT or CD47 KO mice as described above. For the white matter, RNAs from five (WT; 13-15 wks of age) and four (KO; 12-15 wks of age) genotype-matched different male animals were pooled and subjected to the analysis. For the brain mononuclear cells, RNAs from seven (WT; 10-15 wks of age) and six (KO; 12-16 wks of age) male animals were pooled and analysed. Microarray analyses were performed by the Dragon Genomics Center of Takara Bio (Otsu, Japan). The quality of RNA samples was confirmed by an Agilent 2100 Bioanalyzer (Agilent Technologies, Palo Alto, CA). Biotinylated complementary RNA (cRNA) was synthesized using the GeneChip 3‘IVT Express Kit (Affymetrix, Santa Clara, CA) from 250 ng of total RNA prepared from the white matter, or using the Ovation Pico WTA systemV2 (NuGEN, San Carlos, CA) and Encore Biotin Module (NuGEN) from 20 ng of total RNA prepared from the brain mononuclear cells. Following fragmentation, 10 μg of cRNA was hybridised for 16 h at 45°C on the GeneChip Mouse Genome 430 2.0 Array (Affymetrix) with the use of Hybridization, Wash and Stain Kit according to the GeneChip 3’IVT Express Kit User Manual (Affymetrix). GeneChips were then scanned by a GeneChip Scanner 3000 7G (Affymetrix) under the control of Affymetrix GeneChip Command Console Software (Affymetrix). Obtained data were processed using the Expression Console Software (Affymetrix).

#### Transmission Electron Microscopy (TEM)

Sample preparation for TEM analysis was carried out as described previously (Wilke et al., 2013) with minor modifications. Mice were anesthetised as described above and were then perfused transcardially with 1.6% PFA and 3% glutaraldehyde in 0.1 M phosphate buffer (pH 7.4). Brain tissues were removed and fixed with 4% PFA in 0.1 M phosphate buffer overnight at 4°C. Brains were coronally sectioned into slices of 100-μm thickness, and these slices were incubated with a fixative containing 2% reduced OsO4 and 1.5% potassium ferrocyanide in 0.1 M sodium cacodylate buffer (pH 7.4) for 1 h on ice, with 1% thiocarbohydrazide solution for 20 min at room temperature and then with 2% OsO4 solution for 30 min. Sections were incubated in 1% uranyl acetate overnight at 4°C, incubated with Walton’s lead aspartate solution for 75 min at 60°C, dehydrated by sequential treatment for 10 min in each of 50, 70, 80, 90, 95, and 100% ethanol, and then placed in a second solution of 100% ethanol. The sections were incubated with propylene oxide twice, with a 1:1 mixture of propylene oxide/Durcupan resin (Durcupan ACM, Fluka, Buchs, Switzerland) for 10 min, with Durcupan resin twice for 10 min each and then cured on a slide glass at 60°C for 2 days. The cured resins containing the anterior commissure were trimmed out from the flat resins under a stereo microscope and re-embedded into resin blocks for ultrathin sectioning. Ultrathin sections were prepared at a 50-nm thickness using an ultramicrotome (Ultracut-T, Leica) and collected onto single slot copper grids. The samples were examined at ×2500 magnification by a transmission electron microscope (H-7650, Hitachi, Tokyo, Japan) and digital electron micrographs were captured. The frequency of myelinated axons among all axons in the anterior commissure and g-ratios of myelinated axons, the ratio of circumference of axon over myelin, were determined by the use of iTEM software (Olympus SIS, Münster, Germany).

#### Cuprizone model of demyelination

Control (SIRPα^fl/fl^:—) or SIRPα cKO (SIRPα^fl/fl^:Cx3cr1CreER^T2^) mice were fed a 0.2% (w/w) cuprizone [bis(cyclohexanone)oxaldihydrazone, Sigma] (Cpz) diet. Mice were sacrificed and their brains were processed for immunohistochemical analysis after 3 or 5 wks of Cpz treatment. Other groups of mice were returned to a normal diet after 5 wks of Cpz treatment and allowed to recover for 2 wks prior to immunohistochemical analysis. Mice fed a normal diet without Cpz for 7 wks were analysed as controls. Frozen brain coronal sections with a thickness of 20 μm were prepared and stained with specific antibodies for MBP (Myelin Basic Protein), Iba1, CD11c, and Olig2. Coronal slices from each mouse at approximate levels −1.9 to −2.0 mm from the bregma (Paxinos & Franklin, 1997) were analysed by fluorescence microscopy. Acquired digital images were analysed by ImageJ software (Schneider et al., 2012) to quantify the area of demyelination or microgliosis and the number of Olig2^+^ cells.

#### Statistical analysis

Data are presented as the means ± SEM and were analysed by the Student’s t-test. A P value < 0.05 was considered statistically significant.

#### Data availability

The microarray data have been deposited to the Gene Expression Omnibus database (https://www.ncbi.nlm.nih.gov/geo/) (accession numbers: GSE118804 and GSE118805 for the white matter and the brain mononuclear cells, respectively).

## Acknowledgments

We thank E. Urano and T. Maegawa for technical assistance. This work was supported by a Grant-in-Aid for Scientific Research on Innovative Areas (“Brain Environment”), a Grant-in-Aid for Scientific Research (C), and a Grant-in-Aid for Challenging Exploratory Research from the Ministry of Education, Culture, Sports, Science and Technology of Japan, and a grant from Takeda Science Foundation of Japan.

## Author Contributions

H. O. conceived the research and wrote paper; H.O. and M. S.-H. designed and conducted the experiments, and analysed the data; T. I., R. E., and Y. F. performed electron microscopic analysis; T. N., R. T., H. N., A. Horikoshi, M. A., Y. H., W. S., A. Hirose, K. K., and A. S. conducted the experiments and analysed data. T. K., Y. M., Y. S., M. N., K. S., P.-A. O., S. J. and T. M. contributed to the experimental resources and wrote the paper. All authors reviewed the manuscript.

---

**Figure 3—figure supplement 1.**
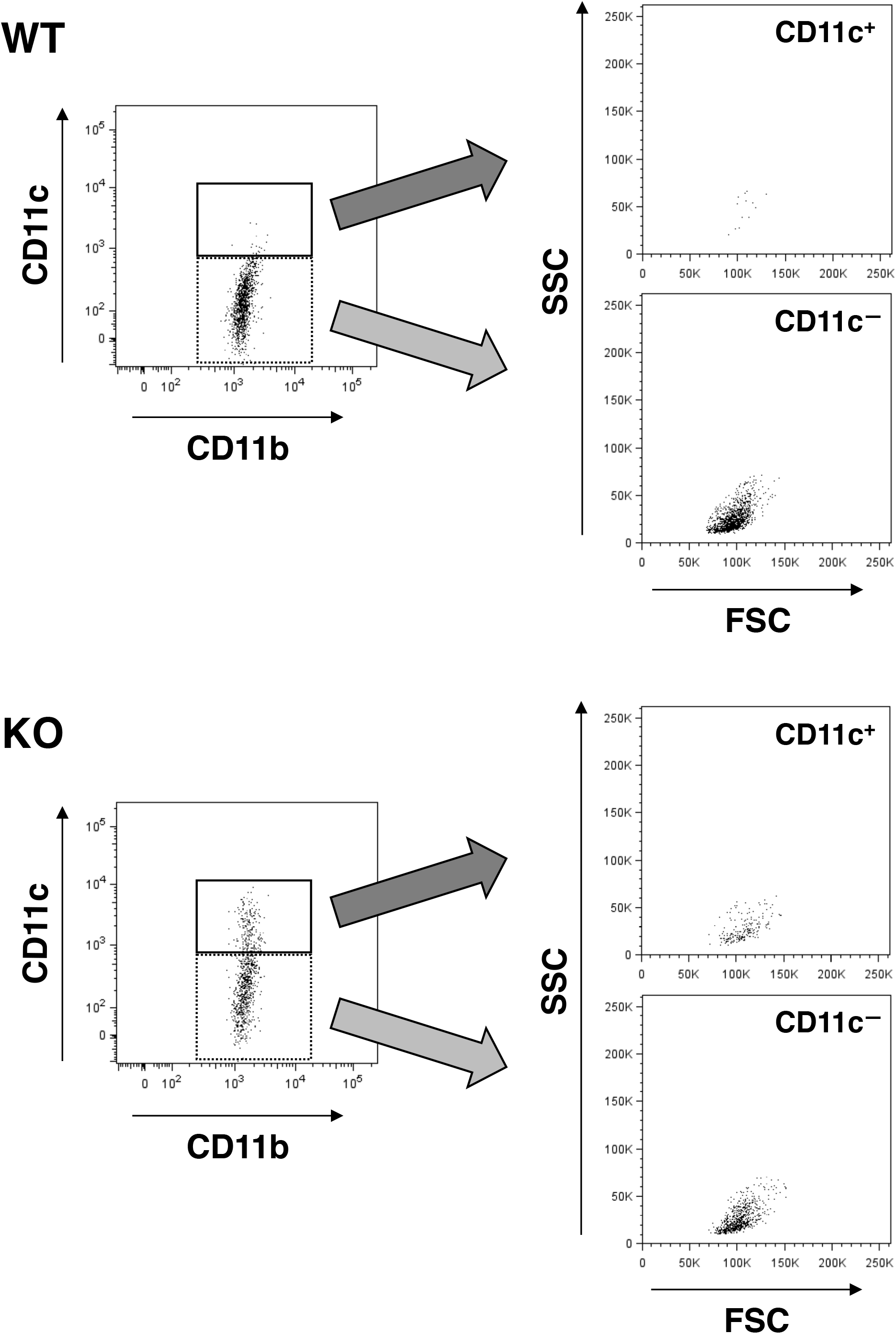
Forward (FSC) and side (SSC) scatter distribution of CD11c^+^ microglia. FSC/SSC distribution of WT (upper panels) and SIRPα KO (lower panels) microglia were analyzed. CD11c^+^ and CD11c— microglia in the left panels (events in the solid and dotted rectangles, respectively) are presented on the FSC/SSC plots in the right panels. Scatter plots in the left panels are identical to those shown in Figure 3B. All data used in Figure 3B were analysed in the same way, and representative results were shown.

**Figure 3—figure supplement 2.**
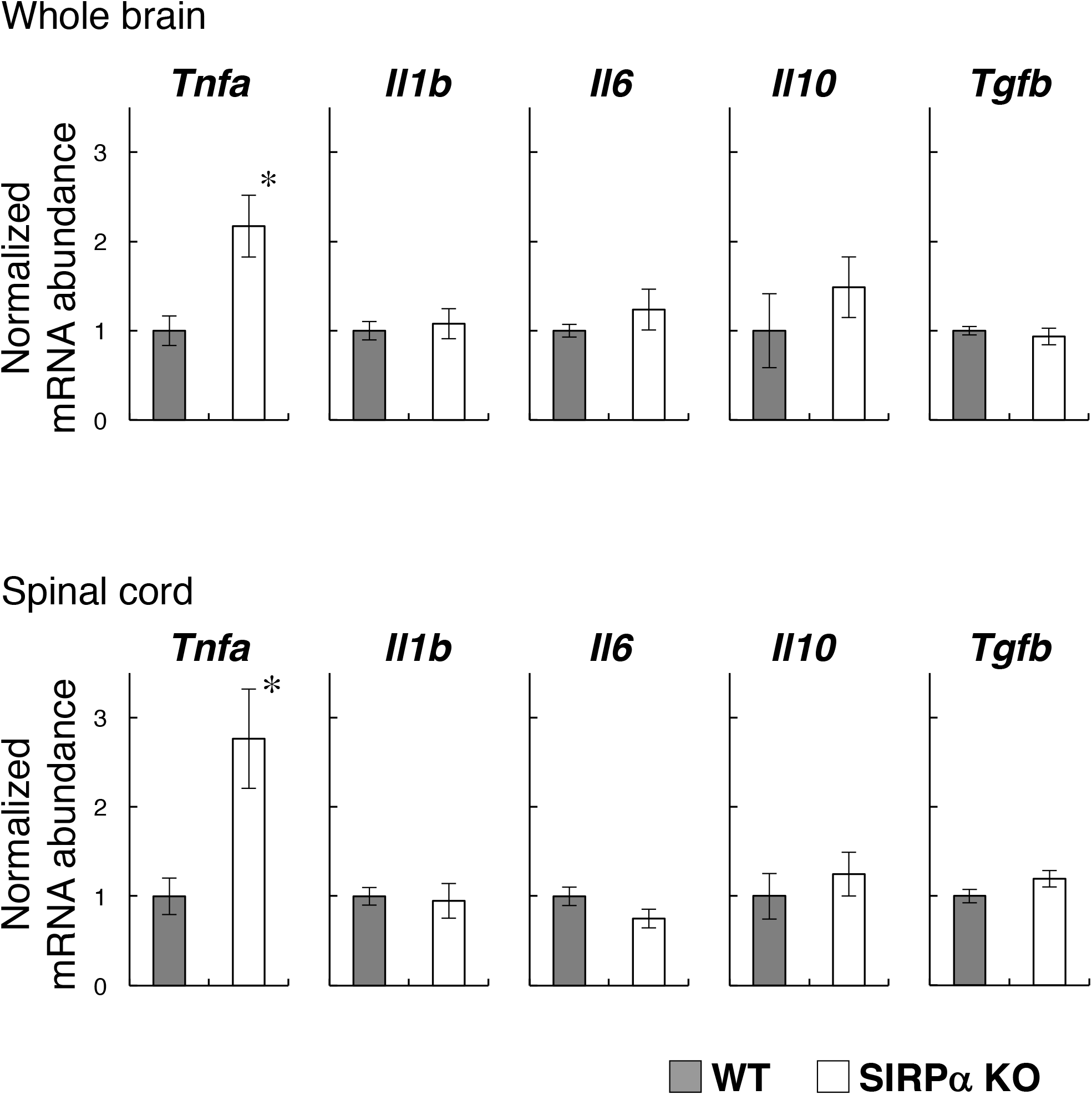
Expression of pro and anti-inflammatory cytokines in the brain and spinal cord of SIRPα KO mice. Whole brains and spinal cords were dissected from WT or SIRPα KO mice at 13–20 wks of age for the isolation of total RNA. The expressions of the indicated cytokines were then determined by quantitative polymerase chain reaction (PCR) analysis. The amount of each mRNA was normalised to that of glyceraldehyde-3-phosphate dehydrogenase (GAPDH) mRNA and is presented relative to the mean value for control WT mice. Data are the means ± SEM (n = 6 mice for each genotype). *P < 0.05 (Student’s t-test).

**Figure 5—figure supplement 1.**
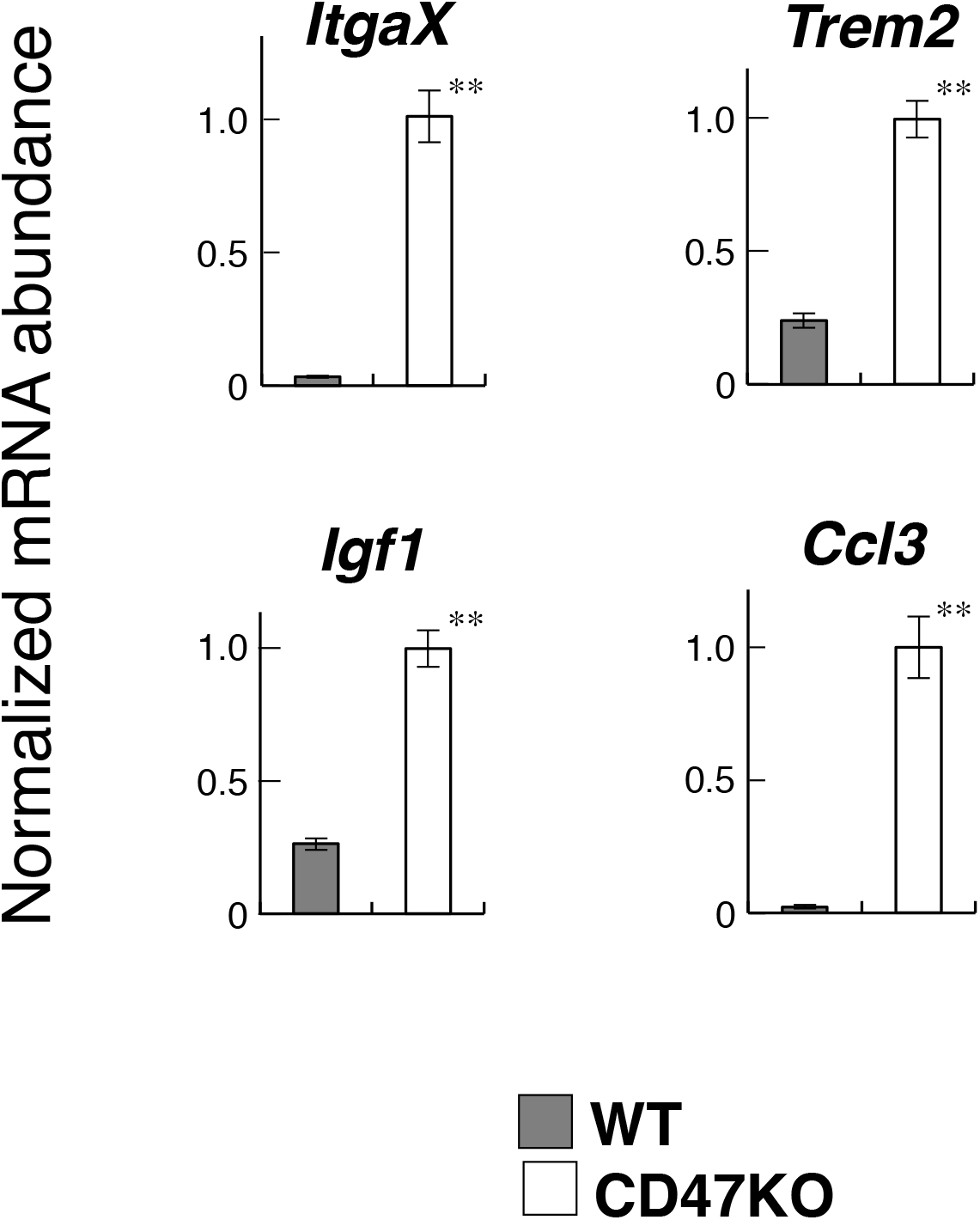
Quantitative PCR analyses of the white matter RNA. Total RNA samples from the optic chiasm and optic nerve of CD47 KO and WT mice (n = 4 and 6, respectively) were subjected to quantitative PCR analysis to determine the expression of the indicated genes. The amount of each mRNA was normalised to that of glyceraldehyde-3-phosphate dehydrogenase (GAPDH) mRNA and presented relative to the mean value for control WT mice. Data are the means ± SEM. **P < 0.01 (Student’s t-test).

**Figure 5—figure supplement 2.**
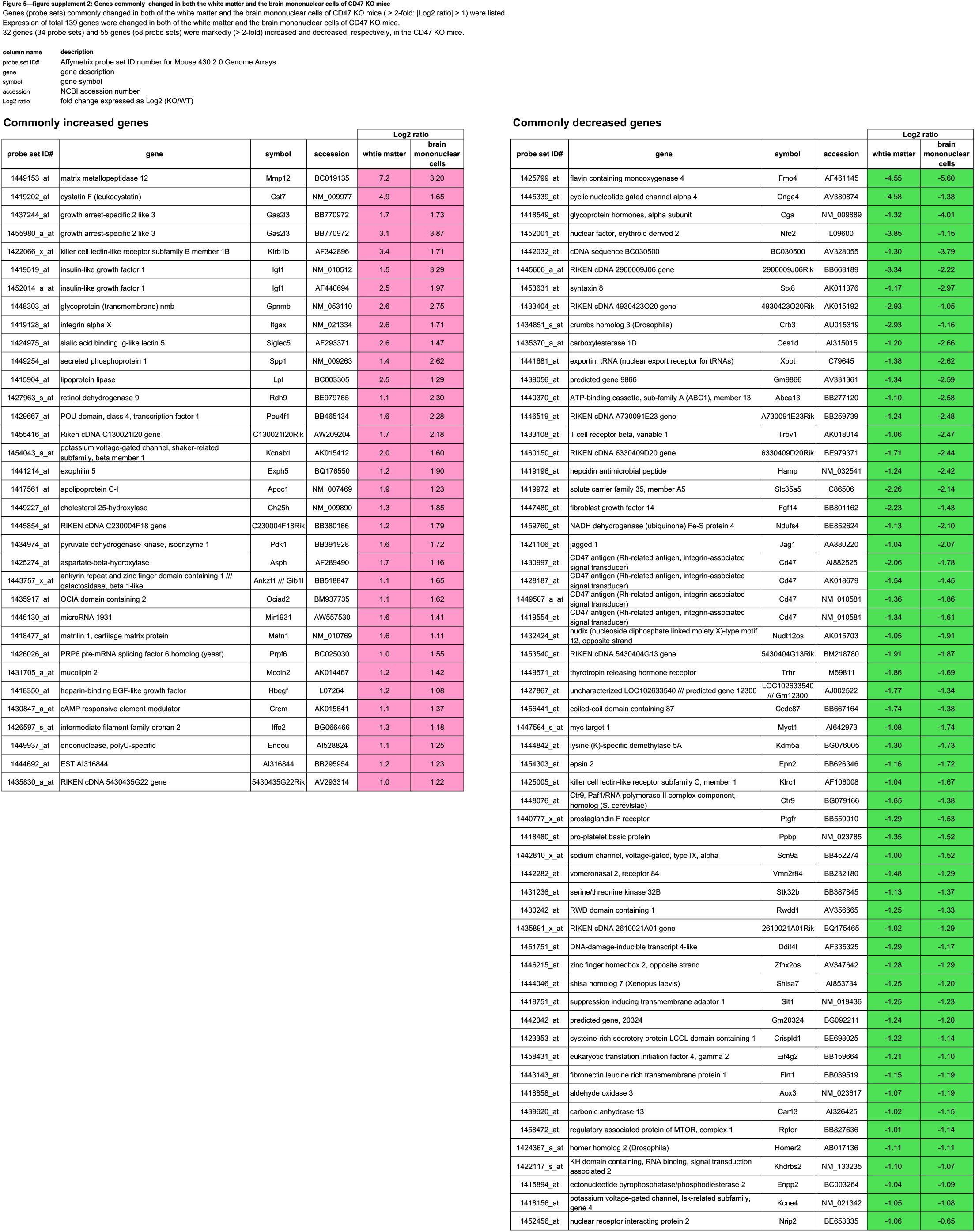
Genes commonly changed in both of the white matter and the brain mononuclear cells. The genes commonly changed more than 2-fold (|Log2 ratio| > 1) in both of the white matter and the brain mononuclear cells of CD47 KO mice were listed. **probe set ID#:** Affymetrix probe set ID number for Mouse 430 2.0 Genome Arrays, **gene:** gene description, **symbol:** gene symbol, **accession:** NCBI accession number, **Log2 ratio:** fold change expressed as Log2 (KO/WT).

**Figure 6—figure supplement 1.**
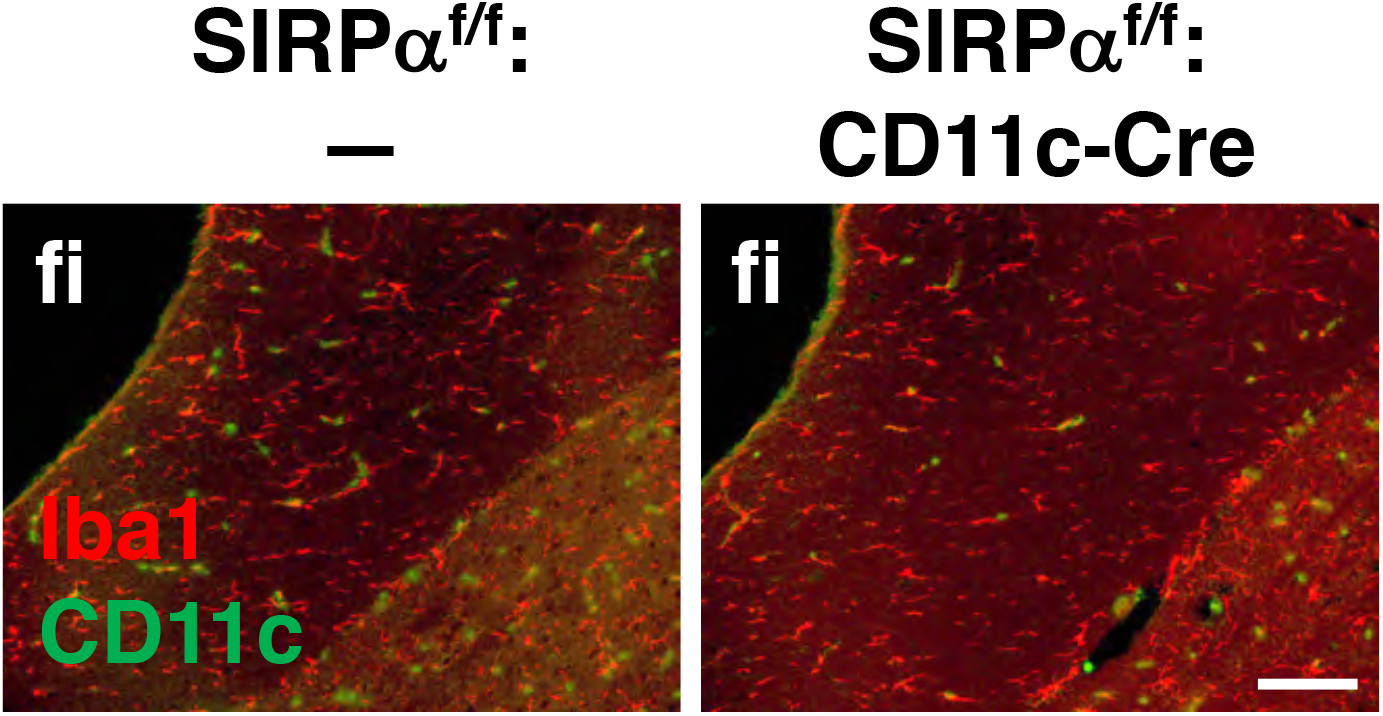
Immunohistochemical analysis of CD11c^+^ cell-specific SIRPα conditional knockout mice. Brain sections prepared from control (SIRPα^fl/fl^:—) and CD11c^+^ cell-specific SIRPα cKO (SIRPα^fl/fl^:CD11c-Cre) mice at 25 wks of age were immunostained as in Figure 6B. Merged images are shown. fi: fimbria. Data are representative of at least 3 independent animals. Scale bar: 100 μm.

Supplementary file 1: Gene expression changes in the white matter of CD47 KO mice. All genes changed more than 2-fold (|Log2 ratio| > 1) in the white matter of CD47 KO mice were listed. **probe set ID#:** Affymetrix probe set ID number for Mouse 430 2.0 Genome Arrays, **gene:** gene description, **symbol:** gene symbol, **accession:** NCBI accession number, **Log2 ratio:** fold change expressed as Log2 (KO/WT).

**Supplementary file 2: Gene expression changes in the white matter of CD47 KO mice**. All genes changed more than 2-fold (|Log2 ratio| > 1) in the brain mononuclear cells of CD47 KO mice were listed. **probe set ID#:** Affymetrix probe set ID number for Mouse 430 2.0 Genome Arrays, **gene:** gene description, **symbol:** gene symbol, **accession:** NCBI accession number, **Log2 ratio:** fold change expressed as Log2 (KO/WT).

## References

Acevedo G., Padala N. K., Ni L., & Jonakait G. M. (2013). Astrocytes inhibit microglial surface expression of dendritic cell-related co-stimulatory molecules through a contact-mediated process. J Neurochem, 125(4), 575–587. doi:10.1111/jnc.12221.

Barclay A. N., & Hatherley D. (2008). The counterbalance theory for evolution and function of paired receptors. Immunity, 29(5), 675–678. doi:10.1016/j.immuni.2008.10.004.

Baumann N., & Pham-Dinh, D. (2001). Biology of oligodendrocyte and myelin in the mammalian central nervous system. Physiol Rev, 81(2), 871–927. doi:10.1152/physrev.2001.81.2.871.

Bulloch K., Miller M. M., Gal-Toth J., Milner T. A., Gottfried-Blackmore A., Waters E. M., … McEwen B. S. (2008). CD11c/EYFP transgene illuminates a discrete network of dendritic cells within the embryonic, neonatal, adult, and injured mouse brain. J Comp Neurol, 508(5), 687–710. doi:10.1002/cne.21668.

Cantoni C., Bollman B., Licastro D., Xie M., Mikesell R., Schmidt R., … Piccio L. (2015). TREM2 regulates microglial cell activation in response to demyelination in vivo. Acta Neuropathol, 129(3), 429–447. doi:10.1007/s00401-015-1388-1.

Caton M. L., Smith-Raska, M.R., & Reizis B. (2007). Notch-RBP-J signaling controls the homeostasis of CD8- dendritic cells in the spleen. J Exp Med, 204(7), 1653–1664. doi:10.1084/jem.20062648.

Chao M. P., Tang C., Pachynski R. K., Chin R., Majeti R., & Weissman I. L. (2011). Extranodal dissemination of non-Hodgkin lymphoma requires CD47 and is inhibited by anti-CD47 antibody therapy. Blood, 118(18), 4890–4901. doi:10.1182/blood-2011-02-338020.

Chiu I. M., Morimoto E. T., Goodarzi H., Liao J. T., O’Keeffe S., Phatnani H. P., … Maniatis T. (2013). A neurodegeneration-specific gene-expression signature of acutely isolated microglia from an amyotrophic lateral sclerosis mouse model. Cell Rep, 4(2), 385–401. doi:10.1016/j.celrep.2013.06.018.

Gitik M., Liraz-Zaltsman S., Oldenborg P. A., Reichert F., & Rotshenker S. (2011). Myelin down-regulates myelin phagocytosis by microglia and macrophages through interactions between CD47 on myelin and SIRPα (signal regulatory protein-α) on phagocytes. J Neuroinflammation, 8, 24. doi:10.1186/1742-2094-8-24.

Goldmann T., Wieghofer P., Müller P. F., Wolf Y., Varol D., Yona S., … Prinz M. (2013). A new type of microglia gene targeting shows TAK1 to be pivotal in CNS autoimmune inflammation. Nat Neurosci, 16(11), 1618–1626. doi:10.1038/nn.3531.

Gudi V., Gingele S., Skripuletz T., & Stangel M. (2014). Glial response during cuprizone-induced de- and remyelination in the CNS: lessons learned. Front Cell Neurosci, 8, 73. doi:10.3389/fncel.2014.00073.

Halleskog C., Mulder J., Dahlström J., Mackie K., Hortobágyi T., Tanila H., … Schulte G. (2011). WNT signaling in activated microglia is proinflammatory. Glia, 59(1), 119–131. doi:10.1002/glia.21081.

Hart A. D., Wyttenbach A., Perry V. H., & Teeling J. L. (2012). Age related changes in microglial phenotype vary between CNS regions: grey versus white matter differences. Brain Behav Immun, 26(5), 754–765. doi:10.1016/j.bbi.2011.11.006.

Holtman I. R., Raj D. D., Miller J. A., Schaafsma W., Yin Z., Brouwer N., … Eggen B. J. (2015). Induction of a common microglia gene expression signature by aging and neurodegenerative conditions: a co-expression meta-analysis. Acta Neuropathol Commun, 3, 31. doi:10.1186/s40478-015-0203-5.

Huang D. W., Sherman B. T., & Lempicki R. A. (2009a). Bioinformatics enrichment tools: paths toward the comprehensive functional analysis of large gene lists. Nucleic Acids Res, 37(1), 1-13. doi:10.1093/nar/gkn923.

Huang D. W., Sherman B. T., & Lempicki R. A. (2009b). Systematic and integrative analysis of large gene lists using DAVID bioinformatics resources. Nat Protoc, 4(1), 44-57. doi:10.1038/nprot.2008.211.

Ishikawa-Sekigami T., Kaneko Y., Okazawa H., Tomizawa T., Okajo J., Saito Y., … Nojima Y. (2006). SHPS-1 promotes the survival of circulating erythrocytes through inhibition of phagocytosis by splenic macrophages. Blood, 107(1), 341–348. doi:10.1182/blood-2005-05-1896.

Jaiswal S., Jamieson C. H., Pang W. W., Park C. Y., Chao M. P., Majeti R., … Weissman I. L. (2009). CD47 is upregulated on circulating hematopoietic stem cells and leukemia cells to avoid phagocytosis. Cell, 138(2), 271–285. doi:10.1016/j.cell.2009.05.046.

Kamphuis W., Kooijman L., Schetters S., Orre M., & Hol E. M. (2016). Transcriptional profiling of CD11c-positive microglia accumulating around amyloid plaques in a mouse model for Alzheimer’s disease. Biochim Biophys Acta, 1862(10), 1847–1860. doi:10.1016/j.bbadis.2016.07.007.

Kaunzner U. W., Miller M. M., Gottfried-Blackmore A., Gal-Toth, J., Felger J. C., McEwen B. S., & Bulloch K. (2012). Accumulation of resident and peripheral dendritic cells in the aging CNS. Neurobiol Aging, 33(4), 681-693.e681. doi:10.1016/j.neurobiolaging.2010.06.007.

Keren-Shaul H., Spinrad A., Weiner A., Matcovitch-Natan O., Dvir-Szternfeld, R., Ulland T. K., … Amit I. (2017). A unique microglia type associated with restricting development of Alzheimer’s disease. Cell, 169(7), 1276-1290.e1217. doi:10.1016/j.cell.2017.05.018.

Kettenmann H., Hanisch U. K., Noda M., & Verkhratsky A. (2011). Physiology of microglia. Physiol Rev, 91(2), 461–553. doi:10.1152/physrev.00011.2010.

Kojima Y., Volkmer J. P., McKenna K., Civelek M., Lusis A. J., Miller C. L., … Leeper N. J. (2016). CD47-blocking antibodies restore phagocytosis and prevent atherosclerosis. Nature, 536(7614), 86–90. doi:10.1038/nature18935.

Kusakari S., Ohnishi H., Jin F. J., Kaneko Y., Murata T., Murata Y., … Matozaki T. (2008). Trans-endocytosis of CD47 and SHPS-1 and its role in regulation of the CD47-SHPS-1 system. J Cell Sci, 121(Pt 8), 1213-1223. doi:10.1242/jcs.025015.

Lampron A., Larochelle A., Laflamme N., Préfontaine P., Plante M. M., Sánchez M. G., … Rivest S. (2015). Inefficient clearance of myelin debris by microglia impairs remyelinating processes. J Exp Med, 212(4), 481–495. doi:10.1084/jem.20141656.

Lindquist R. L., Shakhar G., Dudziak D., Wardemann H., Eisenreich T., Dustin M. L., & Nussenzweig M. C. (2004). Visualizing dendritic cell networks in vivo. Nat Immunol, 5(12), 1243–1250. doi:10.1038/ni1139.

Matozaki T., Murata Y., Okazawa H., & Ohnishi H. (2009). Functions and molecular mechanisms of the CD47-SIRPα signalling pathway. Trends Cell Biol, 19(2), 72–80. doi:10.1016/j.tcb.2008.12.001.

Matsuo A., Akiguchi I., Lee G. C., McGeer E. G., McGeer P. L., & Kimura J. (1998). Myelin degeneration in multiple system atrophy detected by unique antibodies. Am J Pathol, 153(3), 735–744. doi:10.1016/S0002-9440(10)65617-9.

Nishiyama A., Komitova M., Suzuki R., & Zhu X. (2009). Polydendrocytes (NG2 cells): multifunctional cells with lineage plasticity. Nat Rev Neurosci, 10(1), 9–22. doi:10.1038/nrn2495.

Norden D. M., & Godbout J. P. (2013). Review: microglia of the aged brain: primed to be activated and resistant to regulation. Neuropathol Appl Neurobiol, 39(1), 19–34. doi:10.1111/j.1365-2990.2012.01306.x.

Norkute A., Hieble A., Braun A., Johann S., Clarner T., Baumgartner W., … Kipp M. (2009). Cuprizone treatment induces demyelination and astrocytosis in the mouse hippocampus. J Neurosci Res, 87(6), 1343–1355. doi:10.1002/jnr.21946.

Ohnishi H., Murata T., Kusakari S., Hayashi Y., Takao K., Maruyama T., … Matozaki T. (2010). Stress-evoked tyrosine phosphorylation of signal regulatory protein α regulates behavioral immobility in the forced swim test. J Neurosci, 30(31), 10472–10483. doi:10.1523/JNEUROSCI.0257-10.2010.

Olah M., Amor S., Brouwer N., Vinet J., Eggen B., Biber K., & Boddeke H. W. (2012). Identification of a microglia phenotype supportive of remyelination. Glia, 60(2), 306–321. doi:10.1002/glia.21266.

Oldenborg P. A., Zheleznyak A., Fang Y. F., Lagenaur C. F., Gresham H. D., & Lindberg F. P. (2000). Role of CD47 as a marker of self on red blood cells. Science, 288, (5473), 2051–2054.

Paradowska-Gorycka A., & Jurkowska M. (2013). Structure, expression pattern and biological activity of molecular complex TREM-2/DAP12. Hum Immunol, 74(6), 730–737. doi:10.1016/j.humimm.2013.02.003.

Paxinos G., & Franklin K. (1997). The Mouse Brain in Stereotaxic Coordinates. San Diego: Academic Press.

Poliani, P. L., Wang Y., Fontana E., Robinette M. L., Yamanishi Y., Gilfillan S., & Colonna M. (2015). TREM2 sustains microglial expansion during aging and response to demyelination. J Clin Invest, 125(5), 2161–2170. doi:10.1172/JCI77983.

Ransohoff R. M., & Cardona A. E. (2010). The myeloid cells of the central nervous system parenchyma. Nature, 468(7321), 253–262. doi:10.1038/nature09615.

Remington L. T., Babcock A. A., Zehntner S. P., & Owens T. (2007). Microglial recruitment, activation, and proliferation in response to primary demyelination. Am J Pathol, 170(5), 1713–1724. doi:10.2353/ajpath.2007.060783.

Safaiyan S., Kannaiyan N., Snaidero N., Brioschi S., Biber K., Yona S., … Simons M. (2016). Age-related myelin degradation burdens the clearance function of microglia during aging. Nat Neurosci, 19(8), 995–998. doi:10.1038/nn.4325.

Saijo K., & Glass C. K. (2011). Microglial cell origin and phenotypes in health and disease. Nat Rev Immunol, 11(11), 775–787. doi:10.1038/nri3086.

Schneider C. A., Rasband W. S., & Eliceiri K. W. (2012). NIH Image to ImageJ: 25 years of image analysis. Nat Methods, 9, (7), 671–675.

Selvaraju R., Bernasconi L., Losberger C., Graber P., Kadi L., Avellana-Adalid V., … Boschert U. (2004). Osteopontin is upregulated during in vivo demyelination and remyelination and enhances myelin formation in vitro. Mol Cell Neurosci, 25(4), 707–721. doi:10.1016/j.mcn.2003.12.014.

Sierra A., Gottfried-Blackmore, A.C., McEwen B. S., & Bulloch K. (2007). Microglia derived from aging mice exhibit an altered inflammatory profile. Glia, 55(4), 412–424. doi:10.1002/glia.20468.

Wang Y., Cella M., Mallinson K., Ulrich J. D., Young K. L., Robinette M. L., … Colonna M. (2015). TREM2 lipid sensing sustains the microglial response in an Alzheimer’s disease model. Cell, 160(6), 1061–1071. doi:10.1016/j.cell.2015.01.049.

Washio K., Kotani T., Saito Y., Respatika D., Murata Y., Kaneko Y., … Matozaki T. (2015). Dendritic cell SIRPα regulates homeostasis of dendritic cells in lymphoid organs. Genes Cells, 20(6), 451–463. doi:10.1111/gtc.12238.

Wilke S. A., Antonios J. K., Bushong E. A., Badkoobehi A., Malek E., Hwang M., … Ghosh A. (2013). Deconstructing complexity: serial block-face electron microscopic analysis of the hippocampal mossy fiber synapse. J Neurosci, 33(2), 507–522. doi:10.1523/JNEUROSCI.1600-12.2013.

Willingham S. B., Volkmer J. P., Gentles A. J., Sahoo D., Dalerba P., Mitra S. S., … Weissman I. L. (2012). The CD47-signal regulatory protein alpha (SIRPα) interaction is a therapeutic target for human solid tumors. Proc Natl Acad Sci U S A, 109(17), 6662–6667. doi:10.1073/pnas.1121623109.

Wlodarczyk A., Holtman I. R., Krueger M., Yogev N., Bruttger J., Khorooshi R., … Owens T. (2017). A novel microglial subset plays a key role in myelinogenesis in developing brain. EMBO J, 36(22), 3292–3308. doi:10.15252/embj.201696056.

Wolf Y., Yona S., Kim K. W., & Jung S. (2013). Microglia, seen from the CX3CR1 angle. Front Cell Neurosci, 7, 26. doi:10.3389/fncel.2013.00026.

Yona S., Kim K. W., Wolf Y., Mildner A., Varol D., Breker M., … Jung S. (2013). Fate mapping reveals origins and dynamics of monocytes and tissue macrophages under homeostasis. Immunity, 38(1), 79–91. doi:10.1016/j.immuni.2012.12.001.

Zeger M., Popken G., Zhang J., Xuan S., Lu Q. R., Schwab M. H., … Ye P. (2007). Insulin-like growth factor type 1 receptor signaling in the cells of oligodendrocyte lineage is required for normal in vivo oligodendrocyte development and myelination. Glia, 55(4), 400–411. doi:10.1002/glia.20469.

